# Perceptual learning improves discrimination while distorting appearance

**DOI:** 10.1101/2022.09.08.507104

**Authors:** Sarit F.A. Szpiro, Charlie S. Burlingham, Eero P. Simoncelli, Marisa Carrasco

**Author notes:** Joint first authorship (Sarit F.A. Szpiro, Charlie S. Burlingham).

## Abstract

Perceptual sensitivity often improves with training, a phenomenon known as ‘perceptual learning’. Another important perceptual dimension is appearance, the subjective sense of stimulus magnitude. Are training-induced improvements in sensitivity accompanied by more accurate appearance? Here, we examine this question by measuring both discrimination and estimation capabilities for near-horizontal motion perception, before and after training. Human observers who trained on either discrimination or estimation exhibited improvements in discrimination accuracy alongside increased biases in their estimates away from horizontal. To explain this counterintuitive finding, we developed a computational observer model in which perceptual learning arises from increases in the precision of underlying neural representations. For each observer, the fitted model accounted for both discrimination performance and the distribution of estimates, and their changes after training. Our empirical findings and modeling suggest that learning enhances distinctions between categories, a potentially important aspect of real-world perception and perceptual learning.

## Introduction

One of the most remarkable forms of adult brain plasticity is the capacity to develop perceptual expertise. Perceptual learning (PL) is defined as long-lasting improvements in the performance of perceptual tasks following practice (for reviews, see Lu and Dosher, 2022; Sagi, 2011; Watanabe and Sasaki, 2015). PL has been documented in every sensory modality (see review, Seitz and Dinse, 2007), and has been studied using detection and discrimination tasks in behavior (Dosher et al., 2013; Saffell and Matthews, 2003; Szpiro et al., 2014; Tan et al., 2019; Wang et al., 2016; Xiong et al., 2016; Yang et al., 2020), electrophysiology (Crist et al., 2001; Hua et al., 2010; Law and Gold, 2008; Li et al., 2008; Sanayei et al., 2018; Schoups et al., 2001; Van Kerkoerle et al., 2018), neuroimaging (Byers and Serences, 2014; Diaz et al., 2017; Jia et al., 2020; Shibata et al., 2011; Yotsumoto et al., 2008) and computational modeling (Bejjanki et al., 2011; Dosher and Lu, 2017; Sotiropoulos et al., 2018; Wang et al., 2013; Wenliang and Seitz, 2018).

Most studies of perceptual learning have focused on task performance (e.g., improved accuracy for a given stimulus attribute or reduced thresholds in detection and discrimination tasks (Donovan et al., 2020, 2015; Dosher and Lu, 1998; Hung and Carrasco, 2021; Li et al., 2004; Xiao et al., 2008). However, another critical aspect of perception is **appearance**, the subjective perceived magnitude of a stimulus attribute, which can be assessed via estimation tasks. Stimulus appearance is often systematically distorted, relative to physical stimulus properties. For example, although discrimination accuracy is better near cardinal (horizontal or vertical) than near oblique orientations (known as the “oblique effect”), orientations near horizontal appear repulsed away from horizontal, deviating from their physical value (Appelle, 1972; Krukowski et al., 2003; Loffler and Orbach, 2001; Rauber and Treue, 1998). But the effects of PL on appearance, and the relation of these effects to those on sensitivity, remain largely unknown. The conjoint effect of PL on performance and appearance has implications for theories and computational models (Dosher et al., 2013; Stocker and Simoncelli, 2008; Ganguli and Simoncelli, 2014; Wei and Stocker, 2017), neurophysiology (Kamitani and Tong, 2005), and philosophy (Brogaard et al., 2018) of perception and learning. Moreover, this conjoint effect can inform clinical rehabilitation protocols, such as those used for amblyopia (Gu et al., 2020; Levi and Li, 2009; Roberts and Carrasco, 2022) and for developing effective training protocols for acquiring perceptual expertise (e.g., for radiologists, Waite et al., 2019).

The only study to date that has investigated appearance and PL with the oblique effect revealed that training (without feedback) to estimate nearly horizontal motion *increased* repulsive biases away from horizontal, as measured through either an explicit estimate of motion direction or the direction of smooth pursuit eye movements (Szpiro et al., 2014). However, this study did not examine the critical question of how training on a discrimination task — used in nearly all PL studies and rehabilitation protocols — affects estimation biases. Assessing whether and how training on a discrimination task affects repulsive biases would broaden our understanding of how PL affects perceptual representations.

Here, we investigated the behavioral consequences of training with either discrimination or estimation on both tasks. Observers viewed near horizontal motion directions and were initially tested (“pre-training”) on both a discrimination task (indicate “clockwise” or “counterclockwise” direction) and an estimation task (indicate perceived motion direction by adjusting the orientation of a line; **Figure 1A**). All observers exhibited substantial repulsive biases in the estimation task (e.g., for a 4° motion stimulus they indicated a 12° estimate), consistent with previous reports of reference repulsion (Appelle, 1972; Rauber and Treue, 1998). We found that observers also made estimates that were on the other side of the horizontal boundary (e.g., for a +4° stimulus, they indicated a -12° “misclassified” estimate), leading to a bimodal estimate distribution. After three days of training on either task, participants exhibited a similar degree of improvement in discrimination accuracy, as well as a decrease in noise coherence thresholds. Strikingly, training also shifted the estimate distribution further from the horizontal reducing the frequency of misclassified estimates. Thus, PL improved performance at the expense of a more biased stimulus appearance.

**Figure 1.**
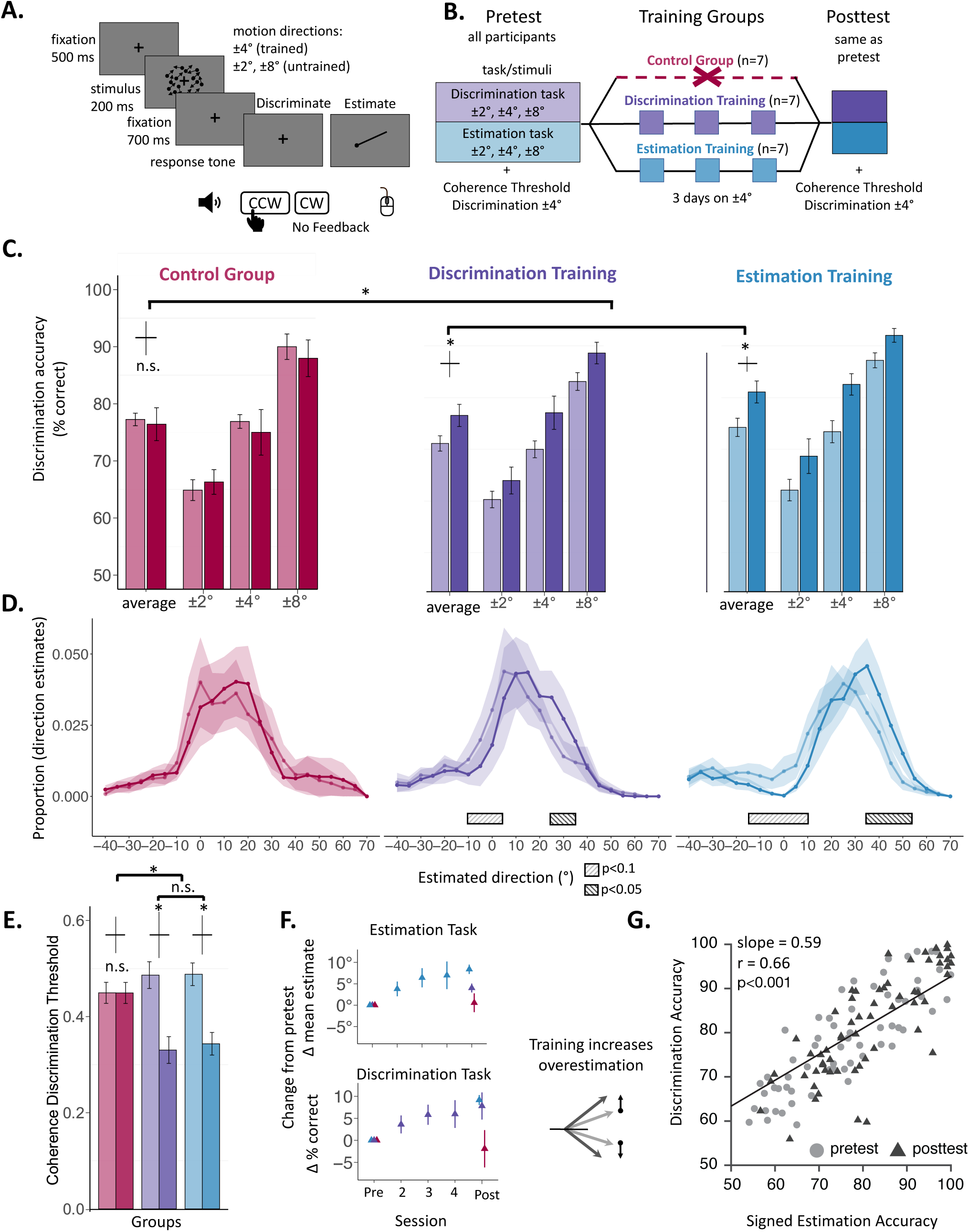
Experimental protocol and behavioral evidence of Perceptual learning. **A**. Trial sequence. **B.** Training procedure for three groups (Control, Discrimination training, Estimation training). **C.** Average discrimination accuracy across groups (indicated with three colors, one per training group) at pre and posttest (unsaturated vs saturated colors, respectively) across directions. Errors bars represent ±1 standard error of the mean with Morey’s correction. **D.** Illustration at the group level of estimate distributions. Mean estimate proportion in each group at pretest versus posttest (unsaturated vs saturated lines, respectively) collapsed across directions and over 5° bins. Shaded area represents standard error of the mean. Significant clusters for the main effect of pre/post training in the cluster mass permutation two-tailed tests are indicated with rectangles (p<0.05 and p<0.1). Note that these estimate plots depict group level estimate distribution for illustration purposes; however, the analysis were based on mixed-linear model that considers individual variability across groups and participants, thereby accounting for the often bi-modal estimate distribution found in individual participants (**Figure 1 – Figure Supplement 1-2, Figure 2)**. **E.** Coherence discrimination threshold for each training group, pretest versus posttest (unsaturated vs saturated lines, respectively). **F.** Performance across days in the estimation and discrimination tasks, for the respective training group. Errors bars represent ±1 standard error of the mean. **G.** Correlation between discrimination accuracy and signed estimation accuracy (percent of direction estimates that were consistent with the correct up/down classification) across all observers in pretest (light circles) and posttest (dark triangles).

To explain the shift in the estimate distribution with PL alongside improvements in discrimination, we developed a computational observer model that makes predictions of discrimination accuracy and the distribution of estimation judgments before and after training. The model is based on three primary assumptions: (1) the internal representation favors cardinal motion directions, which are most common in the natural environment; (2) in the estimation task, observers implicitly categorize motion, and condition their estimates on this choice; and (3) both types of training induce an increase in the precision of representation of trained motions. We find that the simulations of the model, fit to individual observer data, can account for their discrimination accuracy and estimate distribution (including biases and bimodality), both before and after training. Our model provides a link between enhanced sensitivity and perceptual biases. Thus, perceptual learning leads to a more precise representation, which underlies both improved discrimination performance and distorting appearance of stimulus magnitude.

## Results

To examine how PL modifies both discrimination ability and appearance, we asked observers to make judgments about random dot stimuli moving in near-horizontal directions. Observers indicated their responses in either a discrimination task (specifying “clockwise” or “counter-clockwise” relative to rightward horizontal), or an estimation task (indicating the perceived direction of motion by adjusting the orientation of a line). Initially, all observers performed both tasks for motion directions of ±2°, ±4° and ±8° relative to horizontal (**Figure 1A-B**). Following this pre-test, one group (n=7) was trained on the ±4° estimation task, and another group (n=7) was trained on the ±4° discrimination task, for three consecutive days. A third group, the control group (n=7), performed the pre-training tests, but received no additional training over the next three days. Finally, observers from all three groups were tested on both tasks. No feedback was provided in any condition (see Discussion).

All stimuli consisted of a mixture of dots moving coherently in a common direction and a subset moving randomly. Before the experiment, each observer’s noise coherence threshold (the fraction of coherent dots yielding 75% discrimination accuracy) was obtained for the ±4° motion directions using a staircase procedure. This threshold value was subsequently used in testing and training sessions, and the coherence threshold was assessed again after the posttest was completed. In the pretest, neither discrimination accuracy nor estimation judgments interacted with training group for either stimulus motion direction (both F(4,36)<1), demonstrating that prior to training the groups had similar performance in both tasks.

### Discrimination Results

#### Perceptual learning improved discrimination accuracy

For both training groups, PL improved discrimination of motion direction (**Figure 1C**), consistent with prior studies (Saffell and Matthews, 2003; Seitz and Watanabe, 2003). A mixed-design ANOVA revealed a 2-way significant interaction (Training groups vs Control group X Session: F (1,19)=5.79, p=0.02): discrimination accuracy improved significantly between the pre-training and the post-training sessions for observers who trained (Session: F(1,12)=17.35, p=.001), and to a similar degree in both training tasks (Training Task x Session: F (1,12)<1) and across directions (Training Task X Session X Direction: F (2,24)<1), but accuracy was unchanged for observers who did not train (Control group: F(1,6)<1). In sum, discrimination improved regardless of training task and to a similar extent for the trained and untrained motion directions.

#### Perceptual learning decreased noise thresholds

We analyzed noise coherence thresholds before and after training to determine whether there was an improvement in signal-to-noise ratio, as has been found in previous PL research (e.g., (Donovan et al., 2015; Dosher and Lu, 1998; Hung and Carrasco, 2021; Li et al., 2004; Xiao et al., 2008). Due to a technical problem, we do not have the noise thresholds at posttest for three observers, we focus our analysis on the remaining 18 participants. There was a 2-way interaction between group and session (Training groups vs Control group X Session: F(1,16)=8.9, p=0.009; **Figure 1E**). In the control group, without training, coherence thresholds did not change between pretest and posttest sessions, from 47.3% to 44.9%, t(4)=0.04, p=0.99). In contrast, training reduced coherence threshold to a similar degree for both training groups (*Session*: F(1,10)=22.64, p<0.001; *Training Task X Session*: F(1,10)<1): For estimation-training from 48.8% to 34.3% (t (6)=3.55, p=0.01); for discrimination-training, from 47.1% to 33.1% (t(5)=2.97, p=0.03, **Figure 1E**). Across observers, decrements in noise coherence thresholds between posttest and pretest were correlated with increments in discrimination accuracy between posttest and pretest (r=-0.66, p<0.001).

#### Estimation Results

In previous studies researchers have often analyzed changes in estimation judgments by looking at the mean of the estimate distribution (Appelle, 1972; Rauber and Treue, 1998; Szpiro et al., 2014). Here, we found that the bias in the mean estimate increased after training. But we also found that estimate distributions were bimodal (**Figure 1 – Figure Supplement 1**) and the estimate distributions were different between pre and post-test (**Figure 1 – Figure Supplement 2**), so the mean does not provide a reliable characterization of the change in estimates. A change in mean could arise from a change in the locations of the component distributions, or to their relative probability (for an illustration see **Figure 1 – Figure Supplement 3**). In the data, we found both effects – the proportion of estimates that were on the correct side of the discrimination boundary increased (i.e., estimation accuracy) and the correctly classified estimates shifted away from horizontal. We also observed a strong correlation between discrimination accuracy and estimation accuracy. Along with the bimodality, this finding suggests an implicit categorization in the estimation task, and we found including this component in our observer model enabled prediction of estimation behavior.

#### Bimodality in estimate distribution

The distributions of estimates were often distributed bimodally, with a primary mode on the ‘correct’ side of the boundary (i.e., corresponding to the true category of the stimulus, e.g., “CW”), but centered at a biased value (e.g., 15° for observer O4 and directions both ±4° collapsed such that estimates with a positive sign correspond to the correct category; **Figure 2**), and a secondary mode on the ‘incorrect’ side of the boundary (e.g., “CCW”), centered at approximately the negative of the primary mode (e.g., -9° for observer O4 and directions ±4° collapsed such that estimates with a negative sign correspond to the incorrect category; **Figure 2**). In a few cases, there was a third mode centered near the horizontal boundary (**Figure 1 - Figure Supplement 1**). We quantified the shape of these distributions by fitting them with Gaussian Mixture Models containing between one and three components. In the pretest, over half of the estimate distributions were bimodal (63%, over all observers and directions), and the remainder were either unimodal (24%), or tri-modal (13%) (**Figure 1 - Figure Supplement 1**).

**Figure 2.**
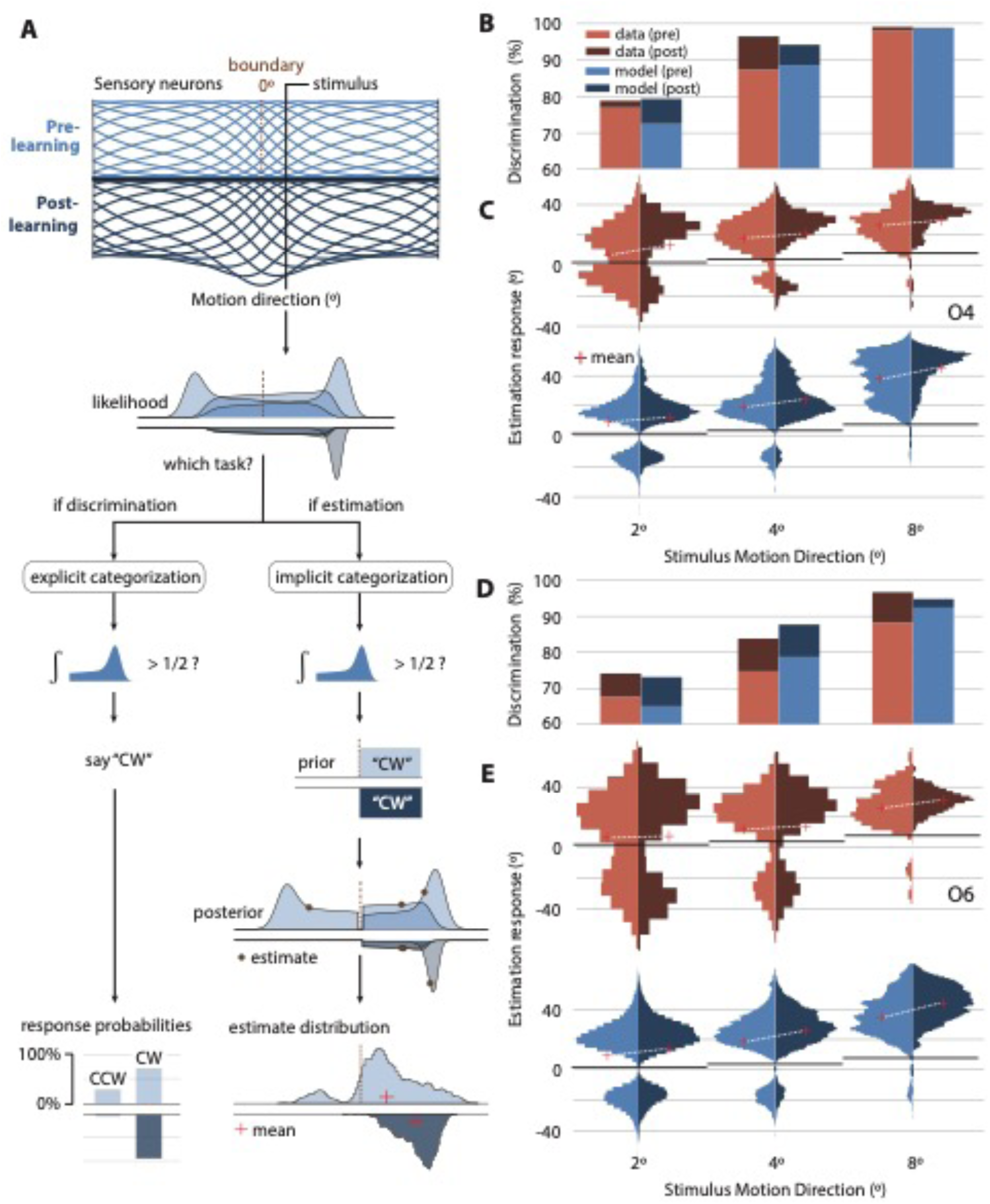
Observer model and human/model behavior for two observers. **(A)** A population of tuned sensory neurons encodes motion direction (tuning curves, representing mean stimulus response; Before training - light blue, upper panels; After training – dark blue, inverted, lower panels). The population is warped such that more neurons represent near-horizontal motion directions, even before training, i.e., via efficient encoding (Ganguli and Simoncelli, 2014; Girshick et al., 2011). Training causes an increase in the gain of the neurons encoding the trained stimuli (±4°), rescaling their tuning curves (inverted, dark blue). For each trial, spike counts for each cell are drawn from independent Poisson distributions, with firing rate governed by the value of a tuning curve at the stimulus’s motion direction. The decoder computes the likelihood of different motion directions given these noisy responses and is assumed to be “unaware” of the fact that motion space is warped. This leads to skewed likelihood functions (3 examples shown). Due to sensory noise, likelihoods fluctuate across trials, with modes sometimes falling on the opposite side of the boundary relative to the stimulus. Regardless of task, the observer performs a discrimination judgement, by comparing the mass of the likelihood on the two sides of the boundary. For the discrimination task, this answer is reported. For the estimation task, this “implicit” discrimination judgement conditions the upcoming estimate (34) — specifically, estimates (3 examples indicated with black points) are computed as the mean of the portion of the likelihood on the side of the boundary corresponding to the chosen category. The mean estimate (red cross) is biased away from the true stimulus by both the efficient encoding (warping) and conditional judgement. And when, by chance, the likelihood falls on the incorrect side of the boundary, its corresponding estimate contributes to a second mode in the estimate distribution, and we label it “misclassified.” The training-induced increase in sensory gain generates fewer likelihoods on the wrong side of the boundary and interacts with the efficient coding/unaware decoding to reduce internal evidence for motion near the horizontal boundary (i.e., “boundary avoidance”). This reduces misclassifications of motion directions (i.e., increases discrimination accuracy) and increases estimation bias. **(B,D)** Comparison of human and model behavior in the discrimination task for two representative observers in the estimation training group (O4, O6). Bar height, discrimination accuracy (% correct) across trials (N =120). **(C,E)** Comparison of human and model behavior in the estimation task. Histograms of estimates (N = 60). Red cross, mean of estimate distribution. White line, change in mean with learning. Black line, stimulus motion direction. Note bimodal shape of estimate distribution (arising from implicit discrimination judgement) and its dependency on learning and stimulus motion direction.

Bimodal distributions, as found in the current study, have been obtained in experiments that require an explicit categorization judgement prior to the estimation response (Jazayeri and Movshon, 2007; Luu and Stocker, 2018; Qiu et al., 2020; Stocker and Simoncelli, 2008). We hypothesized that in the current study observers also performed an implicit categorization (‘CW’ versus ‘CCW’), which influenced their subsequent estimation judgment.

#### Perceptual learning increased the proportion of ‘correct’ estimates

To characterize the change in estimate responses arising from training, we examined the proportion of ‘correct’ estimates - those that fall within the true stimulus category (e.g., “CW” or “CCW”), which we call “signed estimation accuracy’’. Training affected signed estimation accuracy (*Training groups vs. Control group X Session X Direction* F(2,38) p<0.1): Training increased signed estimation accuracy in both training groups, regardless of training task and across directions (*Session*: F(1,12)=25.9, p<0.001; *Session X Direction X Training Task*: F(2,24)<1). In the control group, signed estimation accuracy did not significantly change (*Session*: F(1,6)= 3.45, p=0.112; *Session X Direction*: F(2,12)<1). Thus, in addition to a shift in the distribution, training increased the proportion of estimates on the correct side of the boundary and reduced the probability of misclassifying the stimulus direction in the estimation task.

#### Perceptual learning shifted the estimate distributions away from horizontal

To examine the difference in the distributions of estimation responses between the pretest and posttest, we employed permutation-based cluster mass tests (Maris and Oostenveld, 2007), which identify clusters of adjacent bins of motion directions with a significant difference between pretest and posttest (see **Methods**). It is worth noting that this test does not entail specific assumptions about the underlying shape of the estimate distribution, which could vary among participants (see **Figure 1 – Figure Supplement 1-2,** and results section “The model captures individual differences between observers”). We accounted for individual differences in the distributions using a generalized linear mixed model (GLMM) framework. There was a significant interaction (p<.05) between the training groups and the control group (by permutating group assignments and pre-/post-test session results, see **Methods**). In the training groups – but not in the control group – significant clusters emerged, because in the posttest the estimate distributions differed from the pretest estimate distribution (for the discrimination training the significant cluster was at 25°-35° and for estimation training the cluster was at 35°-55°; **Figure 1D**). Marginally significant clusters emerged where the proportion of estimates at posttest was lower than at pretest (for the discrimination training group the cluster was at -10°–5° and for estimation training group the cluster was at -15°–10°; **Figure 1D**). Importantly, these significant clusters were at each side of the group level estimate distribution, indicative of an overall shift in estimates from horizontal.

The above cluster mass analysis reveals changes that may be driven by either shifts or changes in estimation accuracy (**Figure 1 – Figure Supplement 3**). To examine specific shifts with training only for the correctly classified estimates in the estimate distribution, we conducted an area-under-the-ROC-curve (AUROC) analysis (Locke et al., 2020). The AUROC analysis (see **Methods** for details), the analysis is invariant to changes in estimation accuracy because it examines the overlap between pretest and posttest distributions without considering changes in density. For each direction and observer, we calculated the AUROC between the pretest and posttetst distributions. To examine the magnitudes of these shifts between control and training groups, we conducted a permutation test on the AUROC values and found that the mean magnitude of the shifts was significantly larger in the observers who trained than those who did not (p=0.04, one-tailed test). In sum, training significantly shifted correctly-classified estimates away from horizontal.

To relate our findings to previous studies that have analyzed the mean of the estimates (Appelle, 1972; Rauber and Treue, 1998; Szpiro et al., 2014), we also analyzed central tendency measures of the full distribution. The mean and the median results were larger in the training than in the control groups (see **Appendix 1**). **Figure 1F** illustrates that discrimination and estimation of motion direction changed gradually through the training days. Note that our main statistical analysis and observer model fitting were performed on individual data, rather than on these summary statistics.

#### Strong correlation between discrimination accuracy and signed estimation accuracy

To assess whether observers may have performed an implicit categorization, we examined the proportion of ‘correct’ estimates, the signed estimation accuracy. If observers perform an implicit categorization, they must rely on the same internal representation in both the estimation and discrimination tasks. Therefore, we predicted a correlation between accuracy in the explicit discrimination task and signed estimation accuracy. We found a strong correlation between signed estimation accuracy and discrimination accuracy, collapsed across motion direction and training group for both testing sessions (overall: r=0.66, p<1×10^-6^; pretest: r=0.84; p<1×10^-6^; posttest: r=0.82, p<1×10^-6^, **Figure 1G**), and no significant difference between sessions (Williams test: p=0.766). These results, along with the bi-modal shape of the distribution, support the hypothesis that observers performed an implicit categorization in both tasks, even though they were not instructed to do so. Next, we implemented this assumption in an observer model and show that it allows the model to reproduce the shape of the estimate distributions and their changes with PL.

#### Observer Model

How does PL simultaneously improve discrimination accuracy and modify the estimate distribution? To understand these intriguing findings, we developed an observer model that performs and learns both tasks (**Figure 2A**). The model features three key mechanisms critical for explaining behavior: (1) Neural tuning preferences for cardinal motion, i.e., efficient encoding of motion directions – more resources are devoted to directions that are more common in the natural environment; (2) Conditional inference based on implicit categorization – observers implicitly categorize motion and then condition their estimates on their categorical decision; (3) Gain modulation – increased precision of the internal representation of trained motions in both tasks.

Only gain modulation was assumed to change with training. We analyzed the model’s ability to predict the pre-training behavior and PL effects (differences between the post- and pre-training behavior). In practice, we used the model to simulate either an estimation or discrimination response on each trial of the task, according to the stimulus’s motion direction. And we fit the model to the *entire* distribution of estimates as well as discrimination accuracy, pre- and post-training, minimizing differences between data and model. We found that neural preferences for cardinal motion and implicit categorization allowed the model to explain behavior present even prior to training (estimate bimodality, large estimation biases), and that gain modulation was key for explaining observed PL effects (increases in discrimination accuracy and changes in the estimate distribution).

Specifically, the observer model performs inference based on a probabilistic neural population code (Jazayeri and Movshon, 2007; Ma et al., 2006; Seung and Sompolinsky, 1993). In the encoding stage, a stimulus elicits spikes in a population of motion-direction-sensitive neurons with independent and identically distributed (i.i.d.) Poisson noise, yielding a noisy population response on each trial. The neurons’ tuning curves tile the space of motion directions, but are warped such that they over-represent directions near horizontal (**Figure 2A**, (Fischer and Peña, 2011; Ganguli and Simoncelli, 2010; Girshick et al., 2011), as has been observed in physiological recordings (Xu et al., 2006). This type of inhomogeneity is generally consistent with theories of coding efficiency: more neural resources are devoted to the cardinal directions because they are more commonly encountered in the environment (Ganguli and Simoncelli, 2014). We assumed that the decoding stage is unaware of this warping (Seriès et al., 2009) and therefore wrongly “assumes” that the encoding population is equi-spaced, which leads to perceptual biases away from the horizontal boundary (i.e., “boundary avoidance”). Given that behavioral performance was similar in the two training groups, we assumed that PL arose from an increase in the precision of the internal representation of motion direction, regardless of training group. We modeled this assumption with a gain increase of the sensory neurons encoding the training stimuli (±4°).

The observer model makes either a discrimination or estimation judgement, depending on the task. For discrimination, the observer model reports “counter-clockwise” if most of the internal evidence (likelihood function) lies counter-clockwise of the discrimination boundary, and “clockwise” otherwise. For estimation, the model performs an implicit categorization (using the same discrimination rule), discards internal evidence inconsistent with the chosen category, and reports the mean of the remaining evidence as its estimate ((Stocker and Simoncelli, 2008); see Methods for implementation details). We also assume that the behavioral response itself is noisy in both the estimation task (i.e., additive Gaussian motor noise) and discrimination task (i.e., lapse rate).

#### Model behavior reproduces increased discrimination accuracy and overestimation after PL

For each observer, we fit the model to all behavioral data simultaneously: pre- and post-learning discrimination accuracies and full estimate distributions for all stimuli (±2, ±4 & ±8°). This process yielded parameter estimates shared between the two tasks as well as model behavior for each task separately (see Methods: Parameter estimation).

The model behavior reproduced many aspects of observer behavior, including training-induced increases in discrimination accuracy (**Figure 3A**), overestimation (**Figure 3B**), and signed estimation accuracy (**Figure 3C**), as well as a tight correlation between discrimination accuracy and signed estimation accuracy before and after learning (**Figure 1G**; **Figure 3 – Figure Supplement 1**). Furthermore, the model quantitatively reproduced the transfer of performance from the trained stimuli (±4°) to the tested stimuli (±2 & ±8°; **Figure 2C**, **Figure 3**), including scaling of discrimination accuracy (**Figure 3A**) and reductions in the bimodality of the estimate distribution with learning (**Figure 2C**).

**Figure 3.**
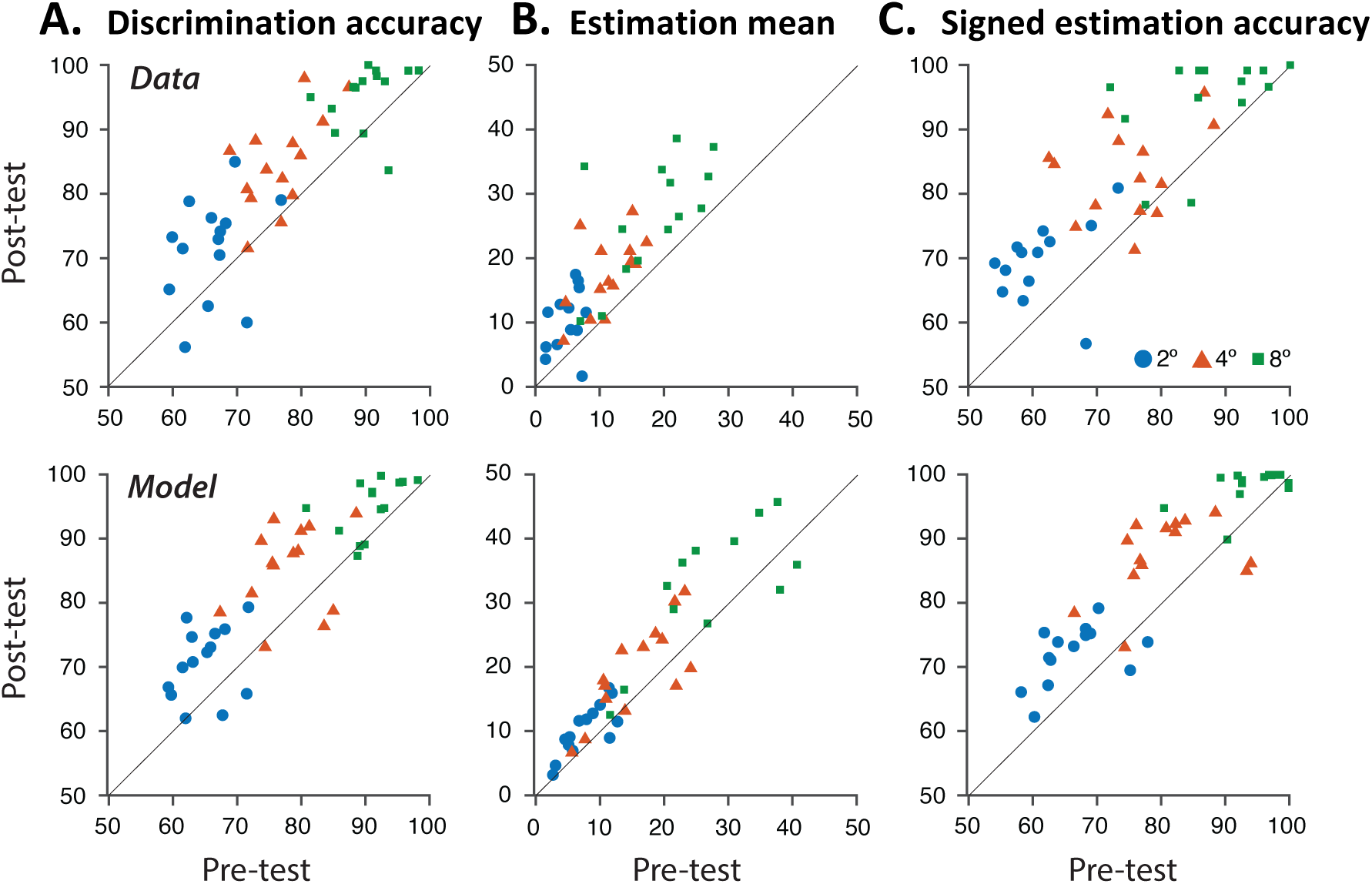
Comparison of human and model performance, over observers trained on both tasks. **(A)** Pre-vs. post-test discrimination accuracy. Top row, human behavioral data. Bottom row, model behavior. Each symbol indicates average discrimination accuracy for one observer for the 2°, 4°, or 8° stimuli (blue circles, red triangles, green squares, respectively). **(B)** Pre-vs. post-test estimation mean. **(C)** Pre-vs. post-test signed estimation accuracy (% of estimates falling on same side of discrimination boundary as the stimulus motion direction). Note similarity between discrimination and signed estimation accuracy (panels A and C) in both data and model.

Specifically, after learning, the model produced fewer estimates that were inconsistent with the true category of the stimulus, leading to a more unimodal estimate distribution, which matched the human observers’ estimate distributions. Across observers, the model explained 85% of the variance in the discrimination accuracy data (p<1×10^-6^), 44% of the variance in estimation mean (p<1×10^-6^), and 52% of the variance in signed estimation accuracy (p<1×10^-6^; **Figure 3**).

#### The model captures individual differences among observers

The model captured large individual differences in behavioral performance, many present even before training. These included nearly unbiased estimation judgements for some observers (e.g., **Figure 2 – Figure Supplement 1**, O12, O15), highly biased estimation judgements for others (e.g., **Figure 2**: O4, O6; **Figure 2 – Figure Supplement 1**: O7), corresponding to fewer or more neurons preferring cardinal motion, respectively (**Figure S10**), as well as low discrimination accuracy (e.g., **Figure 2 – Figure Supplement 1:** O7, O12) and high discrimination accuracy (e.g., **Figure 2**: O4; **Figure 2 – Figure Supplement 1**: O13), corresponding to higher or lower sensory noise, respectively. For observers with little warping / boundary avoidance, corresponding to a more flat environmental prior for motion directions (**Figure 2 – Figure Supplement 1**, O12, O15), the model behavior was similar to existing models of conditional inference (Luu and Stocker, 2018; Stocker and Simoncelli, 2008).

The model also captured individual differences in the effects of PL on behavior, which included notable increases in signed estimation accuracy (**Figure 2**, O4, O6), shifts in the sub-distribution (**Figure 2 – Figure Supplement 1**, O13 vs. O7), and small vs. large increases in discrimination accuracy (**Figure 2 – Figure Supplement 1**, O13 vs. O15). These corresponded mechanistically with decreases in sensory noise with learning, which interacted with individual differences in the sensory noise and cardinal vs. oblique motion neural preferences that each observer came into the experiment with on day 1 (see above).

The model also reproduced the variability in the estimates in the control group (**Figure 3 – Figure Supplement 2**) as well as the clear signs of the implicit categorization decision (i.e., large overestimation pre- and post-training and a tight correlation between estimation and discrimination accuracy).

#### Importance of model components in explaining human behavior

We used model comparisons to qualitatively assess the causal contribution of each model component to the overall behavior (see **Appendix 2**, Reduced Models). In our model, PL arises from an increase in the precision of representation of the training stimuli, which we instantiated by increasing the gain of the training-driven sensory neurons. When we removed this gain modulation (**Figure 2 – Figure Supplement 2**), the model’s behavior was comparable to behavior of observers in the control group (**Figure 3 – Figure Supplement 2**). Note, however, that other mechanisms can produce similar effects. In particular, a model variant in which gain modulation was replaced with tuning changes (such that more neurons in the population encode the trained directions) also captured many features of the data (see Appendix 2; **Figure 2 – Figure Supplement 3**). We focused our analysis on the gain change model, because it provided the best fit to the data, was simpler, and has more support from prior empirical literature (Byers and Serences, 2014; Caras and Sanes, 2017; Hua et al., 2010).

When we removed the efficient encoding component (implemented through warping of the tuning curves), pre- and post-training estimation biases had roughly the right pattern across stimuli and pre-vs. post learning (**Figure 2 – Figure Supplement 4**) but were much smaller than those in the human data. On its own, the efficient encoding step does produce biased estimates (due to the asymmetric posterior), as found in previous studies (Luu and Stocker, 2018; Stocker and Simoncelli, 2008). Yet, the magnitude of this bias is limited by the likelihood function’s width (that arises in our model from a combination of gain, number of neurons, and tuning curve width; Girshick et al., 2011). We concluded that efficient encoding allowed the model to explain the magnitude of overestimation as well as the misclassified estimates observed far from the boundary.

When we removed the conditional inference (implicit categorization) step, the estimate distributions became unimodal — in contrast with bimodal distributions seen in most of the data (**Figure 2 – Figure Supplement 5**). In this model variant, sometimes much of the likelihood function fell on the side of the discrimination boundary inconsistent with the true stimulus category, contributing to the left tail (bottom in figure **Figure 2 – Figure Supplement 5**) of a now unimodal, higher variance estimate distribution (**Figure 2 – Figure Supplement 5**) — instead of contributing to a separate sub-distribution, as when conditional inference is used (**Figure 2E**). Thus, conditional inference allowed the model to explain the bimodal shape of the estimate distribution.

Quantitative model comparisons confirmed our observations about each of the model variants (see **Supplementary Information**). We compared models using two metrics: one was the goodness-of-fit of each model (i.e., the in-sample loss), and the other was the generalization performance (i.e., the out-of-sample cross-validated loss). Each model variant, including the reduced models and the tuning changes models, provided a worse fit of the data than the full model (**Appendix 2– Figure 1A**). The generalization performance of the full model, the no-conditional-inference model, and the tuning-change model (“TC”) were similarly good, with the other models being much worse (**Appendix 2 – Figure 1B**). However, only the full model and tuning-change model (“TC”, which replaced gain changes in the full model with tuning changes) could reproduce the striking bimodal estimation data characteristic of human observers, suggesting that the no-conditional-inference model is inadequate.

## Discussion

The signature characteristic of PL is training-induced performance improvement, most commonly associated with a discrimination task (Ball and Sekuler, 1987; Donovan et al., 2015; Dosher and Lu, 1998; Hung and Carrasco, 2021; Lu and Dosher, 2022; Saffell and Matthews, 2003; Seitz and Watanabe, 2003; Szpiro et al., 2014; Tan et al., 2019; Wang et al., 2016; Yang et al., 2020). We asked how training would concurrently affect stimulus appearance. Whereas one might have expected that improvement in discrimination accuracy would be accompanied by reduced estimation biases (i.e., learning should lead to a more “veridical” representation), our data revealed the opposite. Although participants who trained on either a discrimination or estimation task showed improvement in discrimination accuracy and coherence thresholds, they also exhibited increases in repulsive biases that were present (and substantial) prior to training (**Figure 1D**). Moreover, although training was restricted only to the +4° and -4° directions, these effects were transferred to the untrained nearby directions (+2° and +8°, and -2° and -8°) that correspond to the ‘CW’ and ‘CCW’ categories, illustrating within-category transfer (Szpiro et al., 2014; Tan et al., 2019).

The PL effects we observed bear some similarity to the shorter-timescale effects of attention and adaptation. Specifically, spatial attention has been shown to improve behavioral performance by magnifying task-relevant sensory attributes (Fernández et al., 2022; Ling et al., 2009), even when that entails creating a ‘less veridical’ representation of the stimulus (Mehrpour et al., 2020; review by Carrasco & Barbot, 2019). On slightly longer timescales, adaptation improves direction discrimination for directions near that of the adaptor while repelling perceived directions away from that of the adaptor (Clifford, 2002; Schrater and Simoncelli, 1998). Both these effects are relatively short-lived, and thus do not explain the results of the current study, which increased across multiple days.

Here, we found that the magnitude of bias did not increase over the first training trials, as would have been expected if adaptation mainly modified estimates (see **Appendix 1**). In addition, a previous PL study examined a similar protocol with leftward- or rightwards-motion directions randomly intermixed, thereby preventing adaptation, and found overestimation following training (Szpiro et al., 2014). In the current study, given that direction ‘CW’ and ‘CCW’ motion directions of ±2°, ±4° or ±8° were also randomly intermixed, had adaptation affected estimates, the effect should have ‘canceled’ out across trials (adaptation to ‘CW’ direction would then be followed by adaptation ‘CCW’). Moreover, had observers adapted to the horizontal direction (although never presented), we would have expected smaller estimation biases for motion directions further away from the horizontal (e.g., 8° versus 2°), but that was not the case (see **Figure 3B**). Most critically, the adaptation time scale cannot account for the differences between the training groups and the control group, given that pretest and posttest sessions were identical for all groups. Thus, although short-term effects might have contributed partially to perception, it cannot account for the change in performance across days, the main finding of the study, which is driven by perceptual learning.

As in a number of previous studies, our experiments measure the effects of PL without feedback (Ball and Sekuler, 1987; Guggenmos et al., 2016, 2016; Herzog and Fahle, 1997; Koyama et al., 2004; Petrov et al., 2006). This approach is more consistent with the “unsupervised learning” that occurs in many real-world situations. Moreover, a number of authors have also suggested that it is preferable to avoid feedback when using estimation judgements to assess subjective appearance, in the context of perceptual learning (Szpiro et al., 2014), category learning (Goldstone, 1995), interaction of subjective and objective perceptual organizations (Carrasco and Chang, 1995), 3-D form (Braunstein and Todd, 1990), and appearance of various perceptual dimensions (e.g., (Anton-Erxleben et al., 2013; Carrasco et al., 2004); review by (Carrasco and Barbot, 2019). In discrimination tasks in PL, biased feedback (i.e., reverse incorrect labels of categories) may change the decision criterion, leading to subjective decision biases (Herzog and Fahle, 1999).

PL studies typically use two-alternative choice tasks and only a handful have used estimation tasks. Most of these provided feedback (Green et al., 2015; Grzeczkowski et al., 2019, 2017) and reported reduced estimation variability, but did not examine effects on estimation bias (i.e., appearance). The only study using estimation in the absence of feedback found that estimation training increased overestimation in smooth eye movements and perceptual estimates (Szpiro et al., 2014). Another PL study, which employed feedback during training, used both estimation and discrimination tasks and analyzed cross-task transfer (Green et al., 2015). Training on discrimination had no statistically detectable effect on estimation variance, and training on estimation had no detectable effect on discrimination thresholds. Within-task testing revealed enhanced discrimination performance and reduced estimation variance. These findings suggest that the absence of feedback was critical for our findings of robust cross-task transfer, increased overestimation, and reductions in discrimination thresholds.

How did training lead to changes in the estimate distribution? To explain these findings, we developed a new model of the effects of perceptual learning on discrimination and estimation. We fit the model to the discrimination accuracies as well as the *entire* distribution of estimates, pre- and post-training, meaning we could quantitatively account for non-standard forms of variability within and between participants. Our model combines encoding and decoding components: (1) efficient encoding of cardinal motion directions – more resources are devoted to directions that are more common in the natural environment; (2) conditional inference – observers implicitly categorize motion and then condition their estimates on their category decision; and (3) gain modulation – increased precision of representation of trained motions in both tasks. We show that these components allow the model to explain overestimation and bimodality of estimates pre-training, and the change in these estimates post-training.

The first two model components are needed to explain the pattern of estimate distributions found at pretest (e.g., bimodality of estimate distribution, magnitude of overestimation as well as misclassified estimates far from the boundary). The overestimations we found at pre-training are similar to those in other studies (Jazayeri and Movshon, 2007; Luu and Stocker, 2018; Rauber and Treue, 1998). The first component in the model is efficient encoding that warps the tuning curves around the horizontal, such that more resources are devoted to near horizontal directions because they are more commonly encountered in the environment (Ganguli and Simoncelli, 2014) **Figure 2A**). We show that this model component allows the model to fit the pre-training estimate distributions. This effect and other seemingly anti-Bayesian percepts can be explained by computational models using efficient encoding (Wei and Stocker, 2017, 2015). Although priors are often thought to lead to attractions, in the case of efficient encoding the prior can also lead to repulsions, as in the oblique effect (i.e., high sensitivity and yet high bias near the cardinals). When efficient encoding leads to asymmetric tuning curves with heavier tails on the side of lower prior density, repulsive bias can emerge (**Figure 2A**).

In the current study, the extent of the bias and the bi-modal shape of the distribution suggest that a second component is also in play – implicit categorization. Explicit categorization before estimation judgments has been shown to lead to bias and bimodality in estimate distributions (Jazayeri and Movshon, 2007; Luu and Stocker, 2018; Stocker and Simoncelli, 2008). The implicit categorization component of the model is agnostic as to whether the implicit categorization ‘decision’ is a conscious or unconscious strategy such as observers wanting “to make sure” they are getting the stimulus category correct in the estimation task given the noise in the stimuli. In a prior study with intermixed horizontal and near horizontal directions, there were similar estimation biases for the near horizontal directions (e.g., mean estimates of ∼9° for 3° motion direction (Szpiro et al., 2014). This finding suggests that the estimation biases may emerge even when observers see and estimate horizontal motion and thus are incentivized to not avoid the boundary. Interestingly, implicit categorization, which could also be interpreted as a “post-perceptual” process, interacts with the sensory components, a topic worth investigating in future studies.

Why would observers perform an implicit categorization in the estimation task, when they were not instructed to do so? Empirical (Ding et al., 2017; Zamboni et al., 2016) and theoretical (Qiu et al., 2020; Wu et al., 2009) studies suggest that observers may implicitly commit to a high-level interpretation of a stimulus before estimating it (i.e., conditional inference). This process has been considered as a perceptual analogue of confirmation bias (Nickerson, 1998). Although such biases may seem detrimental for perception, conditional inference may confer distinct advantages (Qiu et al., 2020), including reducing energy costs (Gershman et al., 2015; Simon, 1984), optimizing use of neural resources by discarding unnecessary details about a stimulus (Stocker and Simoncelli, 2008), and protecting crucial information from internally generated noise by storing it in a discrete format (Qiu et al., 2020). Specifically, such a strategy is advantageous when the observer has a good chance of correctly discriminating a stimulus in the presence of post-decisional noise (Luu and Stocker, 2018; Qiu et al., 2020).

Why do observers implicitly categorize around *horizontal*? One explanation is that observers were influenced by the experimental design, either because stimuli were symmetrically distributed around the horizontal boundary or because they were asked to perform the discrimination task during the pretest (albeit in separate blocks of trials). Yet, overestimation has been reported when observers only perform an estimation (Szpiro et al., 2014). Another possibility is that clockwise and counterclockwise of horizontal motion are “natural categories” (Appelle, 1972) shaped by the structure of the world (i.e., cardinal motion is more common than oblique). And because observers exhibit repulsive biases away from cardinals (both clockwise and counter-clockwise), these two groups of percepts clump together over years of experience, forming internal categories which are then used to make perceptual decisions.

A real-world example of this is that of hanging a picture on a wall – very small amounts of tilt are quite noticeable. Indeed, the use of the horizontal as an implicit reference even when it is not explicitly presented has been suggested previously (Dakin et al., 2010). Moreover, the need for a perceptual categorization process to predict anisotropies in motion direction perception, repulsion in estimates from horizontal, *and* high sensitivity near the horizontal has been suggested in computational modelling work (Wong and Price, 2014), although how and at what stage this categorization step takes place is unclear. The emergence of ‘natural’ category boundaries has been recently demonstrated in neural networks trained on objects in natural images, which match with human category boundaries in color perception (de Vries et al., 2022). These examples provide an intuition for the relation between high sensitivity around “anchor” stimuli, and the grouping of nearby values into internal categories.

The third component in the model, gain modulation, is needed to explain how behavioral results changed with training – improved discrimination accuracy and a shift in the estimate distribution away from horizontal. We modeled PL as a change in the gain of sensory neurons representing the trained motion direction. Explanations of the mechanisms underlying PL range from low-level changes in sensory neurons — e.g., changes in gain (Byers and Serences, 2014; Hua et al., 2010) or tuning (Sanayei et al., 2018; Schoups et al., 2001), feed-forward weights, and noise correlations (Bejjanki et al., 2011) — to top-down modulations of neural responses based on context and task (Crist et al., 2001; Li et al., 2004, 2008), changes in decision-making (Law and Gold, 2008), and reweighting (Dosher et al., 2013; Sotiropoulos et al., 2018). Different neuronal changes may underlie performance improvement in PL. A human fMRI study demonstrated that PL may strengthen attentional (gain) modulation of sensory representations in cortex (Byers and Serences, 2014), an account that has been supported by physiological recordings in cat visual cortex (Hua et al., 2010) and gerbil auditory cortex (Caras and Sanes, 2017). Tuning changes in sensory neurons (i.e., sharpening and lateral shifts in individual tuning curves) may also be important for PL (Crist et al., 2001; Li et al., 2004; Sanayei et al., 2018; Schoups et al., 2001). Accordingly, we fit a variant of our model in which we replaced gain modulation with tuning changes (see **Appendix 2**), such that neurons encoding the trained stimuli motion directions would be more densely packed after training (Ganguli and Simoncelli, 2014). This model variant reproduced human behavior nearly as well as the gain-change model, consistent with the fact that various neural mechanisms may explain the observed behavior. Our behavioral data is not sufficient to adjudicate between these possibilities. We speculate that both model variants, changes to sensory neurons in gain or tuning, may potentially account for the decrease in coherence thresholds observed in the data after learning, although we did not model coherence thresholds. Whether gain or tuning, our model and data illustrate that a simple change in sensory neurons may interact with implicit categorization to produce idiosyncratic and unexpected patterns of behavior with learning.

An intriguing aspect of our findings is that PL concurrently reduced noise coherence thresholds and shifted estimate distributions. Previous models and studies of estimation bias (Jazayeri and Movshon, 2007; Luu and Stocker, 2018; Wei and Stocker, 2017, 2015) predict that estimation biases should decrease as sensory noise decreases (e.g., controlled by the motion coherence of an RDK (Jazayeri and Movshon, 2007; Luu and Stocker, 2018). In contrast, in our study, although the noise coherence thresholds *decreased* after training (a known consequence of perceptual learning, e.g., (Donovan et al., 2015; Dosher and Lu, 1998; Hung and Carrasco, 2021; Li et al., 2004; Xiao et al., 2008), estimate distribution shifted from horizontal and mean/median estimates increased. This aspect of the data is not predicted by extant computational models (Luu and Stocker, 2018; Wei and Stocker, 2017, 2015). Our model illustrates how increased precision can account for the coupling between increased estimation biases and increased sensitivity after learning. By linking efficient encoding, conditional inference, and gain or tuning changes, our model suggests that the perceptual system may use different mechanisms in tandem to represent and interpret information more effectively, and better tune to the on-going changes in the environment.

Theories of efficient coding may offer a unifying framework relating PL, discriminability, and perceptual biases. The brain allocates more resources to representing features of the environment that are more common. This efficient allocation of resources must occur through some sort of learning process throughout development — which may well be a form of PL. For example, cardinal orientations (horizontal/vertical) are more common than oblique orientations in natural and retinal images due to the aligning influence of gravity on orientation of environmental structures and visual observers (Girshick et al., 2011). Correspondingly, the brain devotes more sensory neurons to the cardinal orientations, and human observers exhibit better discriminability around the cardinals than obliques, but also larger biases away from them (Appelle, 1972; Krukowski et al., 2003; Loffler and Orbach, 2001; Rauber and Treue, 1998). This is true for many sensory features, including motion direction, and theories of efficient coding can successfully predict discriminability and estimation biases in human and animal behavior based on environmental statistics (Ganguli and Simoncelli, 2014; Girshick et al., 2011; Wei and Stocker, 2017, 2015; Xu et al., 2006).

To account for biases in motion perception that were even present on day 1, the first model component is that the brain devotes more sensory neurons to representing horizontal motion. That is, tuning changes occurred over development due to exposure to non-uniform environmental statistics. On a shorter timescale, across training days, a model variant that assumes a similar mechanism of tuning changes can explain the pattern of results that occurred due to PL. Thus, we suggest that a similar mechanism of efficient coding may underlie PL, whether in development or adulthood, which enhances discriminability at the cost of increasing perceptual biases. Our empirical findings and observer model show that PL-induced increases in the precision of sensory encoding can interact with implicit categorization and a non-uniform internal representation to repel percepts away from the discrimination boundary. Interestingly, repulsion in the appearance of near-boundary stimuli resembles a well-known characteristic of category learning – between-category expansion – the repulsive distortions of values near the category boundary (Goldstone et al., 2001).

Category learning typically enhances discrimination between categories, whereas within-category discrimination is reduced or remains unchanged. For example, in color perception, an object belonging to a group of mostly red objects is judged to be redder than an identically-colored object belonging to another group of mostly violet objects (Goldstone, 1995). That is, color appearance is distorted toward category means, based on the hue statistics of the two groups. Despite some findings of associations between category learning and PL (Green et al., 2015; Tan et al., 2019; Wang et al., 2016), effects of category learning on appearance (e.g., between-category expansion) had not been explored in PL. Our model strengthens the links between category learning and PL (Kattner et al., 2016; Tan et al., 2019; Wang et al., 2016) and demonstrates that distinctions between perceptual categories may be enhanced by changes in sensory encoding (e.g., in gain) over the course of training, without needing to invoke additional changes in cognitive or decision-making processes.

The present findings can have translational implications for real-world manifestations of PL such as perceptual expertise and clinical rehabilitation. For example, when learning to categorize CT images into “cancerous” and “benign” (Dale et al., 2021; Frank et al., 2020; Waite et al., 2019), a radiologist becomes increasingly sensitive to differences between similar images, more accurate, and better at her job. The discriminating features become increasingly salient over training, altering the appearance of both cancerous and benign images such that they appear more dissimilar from each other.

Moreover, our results should also be taken into account when developing protocols and assessment of perceptual rehabilitation of special populations, such as people with amblyopia (Gu et al., 2020; Levi and Li, 2009; Roberts and Carrasco, 2022) and cortical blindness (Cavanaugh et al., 2022, 2019, 2015).

In conclusion, we found that PL improves discrimination and exacerbates estimation biases by shifting the estimate distribution away from the horizontal boundary. To explain these counterintuitive findings, we propose that PL in discrimination tasks may reflect improved categorization, associated with biases in appearance.

## Acknowledgments

This work was supported by a research grant from the NIH RO1 EY016200 to MC, research grant ISF1198/22 to SFAS, funding from the National Institute of Psychobiology in Israel to SFAS, and a National Defense Science and Engineering Graduate fellowship to CSB. We thank members of the Carrasco Lab, especially Shao-Chin Hung and Marc Himmelberg, as well as Mike Landy for useful comments on early versions of the manuscript. We thank Stephanie Badde for helpful comments regarding statistical analysis and permutation tests.

## Methods

### Observers

Twenty-three human adults participated (mean age = 28.9, SD = 1.9; nine males). Experimental procedures were in agreement with the Helsinki declaration and approved by the University Committee on Activities Involving Human Subjects at New York University (FY2016-466). All observers provided written informed consent. All had normal or corrected to normal vision, were untrained and did not know the experiment’s purpose. Two participants did not meet the criterion during pretest to be enrolled in the experiment: One could not perform the estimation task (average estimation deviated more than 70° from veridical); the other could not perform the discrimination task above chance. Thus, twenty-one observers participated in the study. Data is available at https://osf.io/mep9v/?view_only=c97fc183184042c48e45c8a000c793c5.

### Visual stimuli and display

Stimuli were random dot kinematograms (RDK) with dots moving at 15°/s in a stationary aperture with a 5° radius, sparing 0.75° around the central fixation cross. On each frame each dot was assigned a direction that was either the coherent direction or a different random direction according to the coherence level found for each observer, noise dots moved randomly across directions (i.e, directional noise; Brownian motion) and dots were warped when they moved out of the aperture (Seitz and Watanabe, 2003). On each frame 5% of the dots were redrawn to a new position in the aperture to limit tracking of individual dots. Dots were black (4 pixels, 3cd/m2) and were shown on a uniform gray background, with dot density 1.65 dots per square degree. No horizontal reference line was presented to participants. Stimuli were displayed on a calibrated 41×30 cm CRT monitor (IBM P260) with a resolution of 1280×960 pixels, and a 100 Hz refresh rate. Observers were seated at 57 cm distance from the screen with their head supported by a combined chin- and forehead-rest.

### Testing and training tasks

The experiment consisted of five 60-min sessions, one per day, over five consecutive days; the first (pretest) and last (posttest) sessions were identical, and the three intermediate sessions were training sessions (**Figure 1**). The pretest included a short practice on both the estimation and the discrimination tasks. For each observer we tested discrimination coherence thresholds for motion directions of ±4° using three randomly interleaved 60-trial 3-down-1-up staircases. We estimated coherence thresholds by averaging the thresholds reached by the three staircases. This coherence level was then used for all following testing and training.

During pre- and posttest sessions, we measured performance on six randomly presented coherent motion directions (directions relative to horizontal to the right): -8°, -4°, -2° (downwards from horizontal) and 8°, 4°, 2° (upwards from horizontal). Testing sessions (pretest and posttest) included a block of motion discrimination and a block of direction estimation; the order of the two blocks was counterbalanced across observers. Each block consisted of 360 trials. In training sessions, observers were trained either on the estimation task or in the discrimination task, only directions of ±4° directions were presented, and each session consisted of 720 trials presented in four blocks. For the control group, no training was provided, as is the case in some PL studies (Deveau et al., 2014; Kattner et al., 2017; Sabin et al., 2012).

Each trial started with a 500-ms fixation cross at the center of the screen, then the 200-ms RDK appeared followed by a 700-ms ISI after which an auditory start signal indicated that a response could be given (**Figure 1A**). In discrimination blocks, observers pressed a key on the keyboard indicating upward or downward motion relative to horizontal, and the response had to be given within 900 ms. In estimation blocks, a randomly oriented line appeared and, using a mouse, observers were given 4-seconds to orient the line according to the motion direction they perceived; the initial orientation of the line varied uniformly around horizontal with a variance of 5°. Observers mostly responded within the timeline (98% of trials). No feedback was given either for discrimination or estimation tasks as feedback is not necessary for PL to occur and could affect reports rather than appearance (see Discussion).

### Statistical Analysis Methods

For both estimation and discrimination tasks, up and down responses were combined: we negated directions and responses for the downward trials and merged the data with that of the upward trials. We chose this approach also for the estimation task, rather than examining the absolute estimation values, because the negative values of estimations (i.e., to the incorrect side; for a presented direction of 4° a response of -6°) would be considered as relatively ‘correct responses’ (i.e, 6°, or only +2° bias, whereas the original response was -8° bias).

### Mixed Design ANOVA

We used a repeated measures analysis of variance (ANOVA), using session (pretest/posttest) and directions (±2°, ±4° or ±8°) as within-subject factors, and group or training conditions as between-subject factors. When assumptions for sphericity were not met, results were corrected using Greenhouse-Geisser. Mixed design anova was used to analyze coherence thresholds, discrimination accuracy, estimation accuracy, means and modes of the estimate distributions.

### Cluster-mass permutation tests

To examine changes in the estimate distribution without assuming a particular shape for the distribution, we used permutation based cluster mass tests (Maris and Oostenveld, 2007) used to examine significant clusters in previous studies (Balsdon et al., 2020; Ruesseler et al., 2023; Sarasso et al., 2022; Vetter et al., 2019). This allowed us to detect non-parametric shifts in the estimate distributions from pretest to posttest. First, for each participant, estimation data were collapsed across negative and positive directions (-2° were collapsed with the +2°) and across directions (2°, 4°, and 8°). Motion directions were then binned in 5° bins (for pretest and for posttest) between -70° to +70° (0° being horizontal) to create counts of estimates in each motion direction bin for the pretest and posttest. In step 1 of the analysis, for each group we used a Poisson generalized linear mixed model to examine whether the number of estimates significantly differed between pretest and posttest in that motion direction bin across participants in the group (i.e., nEstimates ∼ Session + (1 | subj)). A bin was considered significant if the z-score was above or below the critical z-values in the sampling distribution of the permuted z-scores (see random permutation in step 2; two-tailed, alpha = 0.05). In step 2, we identified clusters of adjacent significant estimate bins with an effect in the same direction (pre>post or post>pre). The clusters that emerged were considered significant by comparing the summed z-score value of the cluster to the distribution of summed z-score values from clusters derived from 1000 random permutations of the data.

To create permuted data, the pre/post labels were swapped within, but not across, participants. Labels were swapped for that participant across all estimate bins. For each permutation, step 1 and 2 were repeated to extract the largest cluster in that permutation. This process led to a distribution of clusters (i.e., of largest summed z-scores across 1000 random permutations). Finally, the original cluster that emerged in data was considered significant if clusters with same or larger values occurred in less than 5% of the randomly permuted datasets (two-tailed values may either be positive or negative depending on whether pretest was larger than posttest or the opposite). This process was done for each training group and a similar process was used to examine significant clusters in the interaction between training groups and control and session (i.e., nEstimates ∼ Group * Session + (1 | subj)). Given that the number of estimates is fixed, a shift between pretest and posttest, would likely mean that if a significant cluster emerged where posttest > prettest, there would likely also be a pretest>posttest cluster in the estimate distribution. Thus, we also illustrated the results for the marginally significant clusters of the randomly permuted datasets but opposite sign (alpha = 0.1, two tailed; **Figure 1D**).

### Permutation based AUROC test

To examine specific changes with learning in the estimate distribution for correctly classified estimates, we conducted a permutation-based AUROC analysis (Locke et al., 2020). For each observer and stimulus motion direction separately, we quantified learning-related shifts (away from horizontal) in the distribution of correctly classified estimates non-parametrically by measuring the area under the ROC curve (AUROC) for the pre- and post-test estimate distributions. This analysis examines the separation between the pretest distribution and the posttest distribution for correctly classified estimates (e.g., a value of 0.5 indicates no separation, whereas a value of 1 indicates complete separation). We obtained p-values for each observer/motion direction pair via permutation tests by shuffling the pre- and post-test distributions together (in 1000 random partitions) and recomputing the AUROC score each time to form a null distribution (one-tailed). All p-values were corrected for multiple comparisons using a procedure for controlling the false discovery rate (FDR) of a family of hypothesis tests (Benjamini et al., 2006).

### Permutation based ANOVA tests for within/between group effects

We used permutation tests to examine central tendency measures of estimation responses in a non-parametric way that is equivalent to mixed design ANOVA. We created 10000 permutations of the dataset. For each permutation the pre/post labels were shuffled within participants for each permutation (e.g., assuming a null hypothesis that there was no difference between pretest and posttest). Then participants were assigned to groups randomly (e.g., again assuming the null that there was no difference between groups) and participants were selected from these random group assignment with replacement to create a random sample of participants in each group in each permutation. For each permutation, the F value of each of the measures of interest was calculated. For example, for the mean estimate we calculated the F value of the interaction between Group X Session with subject as a random factor. Finally, to calculate the p value of the measure of interest (e.g., interaction of Group X Session) in the original dataset we evaluated the percent of F values calculated from the permutations that were equal or larger from the respective F value in the original dataset. This analysis was implemented in R.

### Modeling Methods

We hypothesized that PL in our task can be explained by increases in the precision of the internal sensory representation — a representation that is already warped to devote more resources to certain features. To test this hypothesis, we created a probabilistic observer model that formalizes each of its commitments. Probabilistic observer models (e.g., Bayesian observer models) are most often used to describe how an observer should behave in a task to optimize their performance, given some variability (e.g., uncertainty, ambiguity, sensory noise) that interferes with decision-making (Landy et al., 2011; Shen and Ma, 2019). Such models rely on the assumption that the observer (or some part of their brain) is “aware” of these noise sources, e.g., the distribution of a stimulus’s value across trials, the brain’s noisy measurement of the stimulus, and so on (Seriès et al., 2009). However, it is straightforward to modify the model such that the observer has some set of incomplete or incorrect beliefs (Rahnev and Denison, 2018; Yoo et al., 2021). These “imperfectly optimal observers” (Maloney and Zhang, 2010) can reproduce idiosyncrasies in human behavior while maintaining the fundamental commitment that observers account for their own sensory uncertainty in decision-making rather than ignoring it . Such observer models can account for complexities in human behavior, without being beholden to explaining how each step is part of a normative account or optimal solution. We propose one such observer model, consisting of encoding and decoding stages, which “learns” and performs both the discrimination and estimation tasks. The code for the computational model will be available online: https://github.com/csb0.

### Task Structure

On every trial, there is a 0.5 probability of the stimulus motion direction being CW of horizontal, expressed as *p(C = CW)* = 0.5, where *C*, the category of the stimulus, takes the value CCW or CW. The overall stimulus distribution, *p(s),* is the normalized sum of six delta functions at ±2, ±4, and ±8°, i.e., the probability of each of the six possible motion directions is equal. Once the category is drawn on each trial, the new, “category-conditioned” stimulus distribution *p(s|C)* is the half of *p(s)* on the side of the discrimination boundary (0°) that is consistent with the category *C*.

### Encoding

We assume the stimulus is measured (“encoded”) by a population of *N* = 10 neurons whose tuning curves tile the space of motion directions but are warped such that more neurons represent motion directions near the horizontal, consistent with physiological measurements and theoretical studies of efficient encoding (Ganguli and Simoncelli, 2014, 2010). The tuning curves specify each neuron’s mean firing rate as a function of stimulus direction. Each tuning curve, before warping, is a single cycle of a cos^2 function raised to a power (0.3745) such that its full-width at half maximum (FWHM), *w_t_*, is 71° (Ganguli and Simoncelli, 2014), consistent with typical tuning of motion-direction-selective neurons in macaque area MT (Albright, 1984). Each neuron has a baseline firing rate, *b*, and a gain, *g,* which specifies the above-baseline firing rate for the preferred stimulus (i.e., the maximum response). The pre-training gains, *g_pre_*, are assumed to be identical across the population. The post-training gains are elevated according to a Gaussian profile across the population, centered at 0°, with a standard deviation equal to the spacing between neurons (specifically, with ten neurons, the standard deviation is 4°). A parameter *g_post_* specified the gain of the most responsive neuron in the population. This captures our assumption that learning modulates the gain of sensory neurons, relative to their sensitivity to the trained motion directions.

Warping in our model embodies the efficient coding notion of devoting more neurons and spikes to more common stimuli (Ganguli and Simoncelli, 2014, 2010). This leads to a behavior that resembles avoidance of the boundary when estimating the stimulus, so we refer to this as the efficient encoding / “boundary avoidance” component of the model. The neural population is formed from a set homogeneous (“convolutional”) set of tuning curves that are re-mapped by warping the motion direction axis according to a parametric function that is fit to the data of each observer, but assumed unchanged throughout the experiment (i.e., unaffected by learning). The warping function is defined as the cumulative integral of a cell “density” function: *h(s,a,w_b_,σ_b_) = a* + 2 + ɸ(*s,-w_b_ /2,σ_b_*) *+* ɸ(*-s,w_b_ /2,σ_b_)*, where ɸ is the cumulative Gaussian function. Motion direction is designated with variable *s*, and the shape of the function is governed by three parameters: an amplitude, *a*, that controls the overall magnitude of the warping (i.e., how “extremely” the observer avoids the boundary); *w_b_*, the width of the region over which tuning curves are drawn toward the boundary; and *σ_b_*, the standard deviation of the cumulative Gaussians, which controls the fuzziness of the template’s edges, leading to warping that is more graded or discrete. Examples are shown in **Figure 2 – Figure Supplement 7.** Before usage, the function *h* is numerically normalized to integrate to 1.

For a stimulus direction *s*, each neuron’s spike count *r_i_*, is drawn independently from a Poisson distribution, with rate determined by its tuning curve evaluated at the (warped) value of *s*. The noisy “population response” (i.e., the vector of spike counts from each neuron), denoted **r**, is a sample of the “measurement distribution” *p(**r**|s)*.

### Decoding

The internal representation of the stimulus **r** must be “read out” or decoded to discriminate or estimate motion directions. For this purpose, we used a *maximum a posteriori* or MAP decision rule for the discrimination judgment, and Bayes-least-squares rule for the estimation task. Our methods are identical to those reported in other studies (Jazayeri and Movshon, 2006; Ma et al., 2006) so we refer the reader to their methods for details, and simply provide an intuition here.

Both decoders rely on a likelihood function, in which the measurement distribution is expressed as a function of the stimulus, *s*, for each noisy population response ***r.*** For our model, the log likelihood is equal to the sum (across neurons) of the log of each tuning curve, each weighted by the observed spike count of the associated neuron (Jazayeri and Movshon, 2006; Ma et al., 2006). We incorporate one additional assumption, that the decoder is ‘unaware’ of the warping in the encoder, and so assumes a homogeneous (“convolutional”) encoding population in computing the likelihoods. This idea has been used successfully in the domain of adaptation to explain biases in perception (Seriès et al., 2009).

The likelihood fluctuates randomly on each trial, due to the variability of **r**. In the case when *w_b_* or *a* is zero, the likelihood is symmetric and centered on the true stimulus motion direction, i.e., its mean is an unbiased estimator of the true stimulus (equivalent to probabilistic population coding; (Jazayeri and Movshon, 2006; Ma et al., 2006). Its width, which corresponds with the observer’s uncertainty about the motion direction generating the internal measurement **r**, depends on gain and baseline firing rate. If *w_b_* and *a* are larger than zero, the likelihood is stretched away symmetrically in half around the boundary a distance depending on *w_b_*, and suppressed near the boundary an amount depending on *a.* The likelihood is therefore asymmetric, and the shape of its tails depends on *σ_b_*.

In the discrimination task, the decision variable *d* represents the log posterior ratio, and is determined by the proportion of the likelihood function that falls on each side of the discrimination boundary: If most of the likelihood mass falls on the positive side of the boundary, *d* > 0, the observer reports “CW” (and vice versa). The distribution of discrimination responses across trials is *p(C_est_|C)* and the discrimination accuracy *p(Correct)* = *p(d>0|C=CW)/2* + *p(d<0|C=CCW)/2*. *C_est_* denotes the estimated category. Note that *d* inherits its variability from the population response **r**. To model lapses in the discrimination judgement, we multiply *p(Correct)* by the same factor *1-λ* for all stimulus motion directions. Note that, in our model fitting, *λ* is never allowed to go below 0.5.

In the estimation task, the estimate is computed as the expected value of the posterior distribution, i.e., the Bayes-least-squares estimate (Stocker and Simoncelli, 2008). The posterior is computed on each trial by imposing an implicit discrimination judgment using the rule described above, using the response (CW or CCW) as a conditional prior, *p(s|C_est_)*, which has a value of 1 on the side of the discrimination boundary corresponding to the chosen category, and 0 on the other side (i.e., uniform across the chosen category of motion directions). The product of this conditional prior *p(s|C_est_)*, and the likelihood *p(**r**|s)*, is the posterior *p(s|**r**,C_est_)*, and the estimation response on a single trial is the mean of this posterior. The distribution of estimates across trials *p(s_est_|s)* is computed numerically for each stimulus, ±2, 4, and 8°, via Monte Carlo simulation. To model motor noise in the execution of the response (orienting an arrow with a mouse), i.i.d. Gaussian noise with standard deviation *σ_m_* is added to each estimate distribution.

### Parameter Estimation

The model has six parameters that are fit to individual observers: *a, w_b_, σ_b_, g_pre_, g_post_, λ,* and five parameters that are shared across observers *b, w_t_, N, σ_m_, and σ_g_*. We estimated the individual parameters by minimizing a loss function L(**ϴ**) (with **ϴ** a vector containing the parameters) expressing the fit between each observer’s data and model behavior. L(**ϴ**) was defined as the weighted sum of two terms: (1) absolute difference (L1 norm) between the discrimination accuracy in the data vs. model behavior (i.e., sum of L1 differences across stimuli and pre-vs. post-learning); and (2) the energy distance (Sejdinovic et al., 2013) between the estimate distributions for the data vs. model (sum of distances across stimuli and pre-vs. post-learning). This distance metric takes into account the entire shape of the estimate distribution and is appropriate for comparing distributions with complex shapes (i.e., like the bimodal distributions we observed). We computed the energy distance as implemented in the Information Theoretic Estimators Toolbox in Matlab (Szabó, 2014). The weights on the two terms were used to rescale the two terms in the loss function into a similar numerical range. These weights were computed by evaluating the objective function on an initial random set of parameters, as is common practice in multi-objective optimization (Marler and Arora, 2004). The loss function was stochastic, varying between each run of simulated trials due to sampling variability in the Poisson spike generation. We set the number of simulated trials to 1500 and repeated the optimization 10 times with a new set of initial random parameters.

We fit all free parameters simultaneously for each observer separately using an optimization algorithm designed for stochastic objective functions (Acerbi and Ma, 2017). We ran it 10 times per observer, chose the iteration with the lowest loss. The resulting best-fit parameters were then used to evaluate the model using 1500 simulated trials, generating (stochastic) model behavior for the estimation and discrimination tasks (**Figure 2B,C**, **Figure 3**).

Note that, for each observer, we fit the 6 free parameters with approximately 486 data points (80 estimation responses x 2 pre/post-training x 3 motion directions, plus 1 discrimination accuracy x 2 pre/post-training x 3 motion directions), so this is a well-constrained optimization problem (as compared to fitting only the estimation biases for the 6 stimulus directions).

### Model Comparison

We compared models using iterated *k*-fold (*k* = 3) cross-validated loss. For each observer and each model, we averaged the CV-loss across folds and iterations (folds x iterations = 102), subtracted this from the average CV-loss from the full model, and then averaged this across observers to quantify each model’s generalizability relative to the full model (**Appendix 2 - Figure 1**). We also did the same with the average in-sample (training set) loss across observers to quantify goodness-of-fit relative to the full model. To perform the cross-validated model fitting, for each observer, we computed one normalization factor for each objective (estimation and discrimination) based on a fixed initial set of parameters and kept these normalization factors fixed across models. Note that these factors were quite similar across observers, but not identical. This was essential to keep the training and test losses in the same space across models and hence comparable.

## Appendix 1

### Reaction times

Although our main measures were estimation performance and discrimination accuracy, we also analyzed reaction time (RT) on correct trials and found that there were no speed-accuracy tradeoffs, reaction time decreased between pretest and posttest sessions (Estimation: F (1,18)=12.88, p=.002; Discrimination: F (1,18)=6.16,p=.02) but not differentially between training groups (interaction session X direction X training task, Estimation: F (4,36)=1.77, p=.16; Discrimination: F (4,36)<1).

### Perceptual learning effects on central tendency measures of the estimate distribution

To examine the effect of perceptual learning on the estimate distributions, in addition to analyzing the shift in the estimate distribution between pre- and post-test, we analyzed performance in the estimation task in terms of the mean, median, and mode of the estimate distribution, which are suited for characterizing bi-modal distributions. To analyze these measures, we conducted permutation-based ANOVA tests to examine interactions and main effects compared to the null hypothesis (see **Methods)**. To preview, analyses indicate that the mean and medians of the estimate distributions are shifted further away from horizontal after training. Thus, whereas the mode, the most frequently reported estimate bin did not change, the central tendency measures that capture the full distributions, the mean and the median, shifted away from the pretest after training.

In terms of the fractions of central tendency values that changed after training: For the mean, the means increased in 21, 19, versus 12 cases in estimation and discrimination training groups versus the control group (respectively by group), of the total 21 directions/ subject combinations in each group (7 subject by 3 directions). For the median, for estimation and discrimination training groups versus the control group, the modes increased in 21, 18, versus 15 cases (respectively by group) of the total 21 directions/ subject combinations for each group. For the mode, for estimation, discrimination training groups and the control group, the modes increased in 15, 12, and 12 cases (respectively by group) of the total 21 directions/ subject combinations.

#### A. Estimate means

Previous studies have demonstrated substantial repulsive biases in perceived motion directions near the cardinal axes (Appelle, 1972; Rauber and Treue, 1998; Szpiro et al., 2014). Consistent with this repulsive bias, our observers estimated motion directions near horizontal as being further away from the horizontal boundary: On average, ±2° motion was estimated as ≈±4.6°, ±4° as ≈±11.8°, and ±8° as ≈±19.1°.

Importantly, training improved discrimination accuracy but increased estimates’ mean further away from the horizontal. There was a significant two-way interaction in the average estimate between observers in the training and control groups (*Groups X Session*, p= 0.0357, permuted null, see **Methods**). In the control group, mean estimates did not change (Session: p=0.24). In the training groups there was an *increase* in estimates’ mean (Session: p= 0.05). For example, directions of ±4°, which were perceived as ≈±12° in the pretest, were perceived as ≈±17.85° in the posttest (**Figure 1F**). For both training tasks, estimates’ mean increased to a similar extent for trained and untrained motion directions (*Training Task X Session X Direction*: p= 0.88). There was no interaction across groups (*Training task X Session*: p=0.35) and there was a main effect of session (p= 0.05). In sum, for both training groups, training modifies appearance by increasing the mean estimates away from the horizontal for both the trained and untrained directions. Notably, the increase in mean estimates between pretest and posttest was correlated with the decrease of noise coherence thresholds across observers (R=-0.49, p<0.01).

#### B. Estimate medians

We examined the median, which splits the distributions into equal halves and is less influenced by the precise shape of the distribution and its outliers (**Figure 1**, **Figure 1-Figure Supplement 1-2).** There was a marginal two-way interaction (*Groups X Session*: p=0.086). In the control group, the median did not significantly change (*Session*: p=0.47). Whereas for the training groups, the median increased with training (*Session*: p= 0.007) and there were no interactions with training task or direction (*Training Task X Session*: p= 0.29; *Training Task X Session X Direction*: p= 0.38). Thus, training shifted the median of the estimate distributions further away from the horizontal.

#### C. Estimate Modes

The median and mean are representative estimates of the distribution, and more appropriate than the mode, which only captures the most frequent value and may not necessarily change even though training shifted the distribution away from horizontal. To analyze the modes, we chose the most frequently reported bin in each estimate distribution (as used in the cluster mass test, see **Figure 1D**). Indeed, unlike the median and the mean, at the group-level the mode was not significantly altered by training as examined using permutation tests (*Groups X Session:* p=0.31). Nevertheless, we did note a small but detectable mode shift for a subset of participants (**Figure 1- Figure Supplement 1-2, Figure 2 – Figure Supplement 2**).

### Ruling out adaptation: Estimation responses did not systematically increase over the first few trials of the training

To rule out the possibility that adaptation accounts for the large estimation biases we observed, we analyzed the first 50 trials of the training for the estimation task. If adaptation underlies the large estimation biases we observed, absolute estimation biases should increase over the course of the first few trials of the experiment. For this analysis, we define estimation bias as the difference between the estimate response and the value of the stimulus presented (e.g., for a stimulus value of 2° and an estimation response of 5° the estimation bias would be 3°). To test this possibility of adaptation, we fit linear regression models to the first 50 trials of the training (session 2) of the group that trained on the estimation task (n = 7) and the trained stimulus direction (-4° and +4°). We found that the mean absolute best-fit slopes (grand mean, 0.26°) — i.e., the change in absolute estimation response per trial —did not differ from flat slopes expected under the null hypothesis (grand mean, 0.005°). That is, we could not reject the null hypothesis that the slopes we observed were greater than what would be expected from sampling error alone. We obtained this result using 14 one-tailed permutation tests (14 tests – for n=7 participants and 2 directions per participant), wherein for each stimulus and observer, we shuffled the first 50 estimates for that stimulus and then computed a null slope, repeated this 1000 times, and compared the observed best-fit slope to the null distribution. 0/14 tests were significant with a corrected alpha of 0.0036 following Bonferroni correction. This demonstrates that the estimation responses did not increase with training due to adaptation.

Note that the stimuli (-4° and +4°,) were randomly interleaved across training trials, which would presumably prevent neural adaptation at the timescale of individual trials (as is typically used in experimental manipulations of adaptation). Also, the mean absolute estimation bias on the very first trial in the experiment across participants was large - a bias away from ±2°of 1.8° (i.e., mean estimate of 3.8°), from ±4 of 7.7° (i.e., mean estimate of 11.7°), and from ±8° of 8.8° (i.e., mean estimate of 16.8°). This means that participants came into day 1 of the experiment already with biases, consistent with our modelling assumption that the biases arise in part from observers’ individual neural tuning for motion direction.

### Appendix 2

#### Reduced models

To evaluate the effects of the primary model components (i.e., gain modulation, conditional inference, and efficient coding), we created three reduced models by removing model components one-by-one (**Figure 2 – Figure Supplements 2-4**): (a) For the no-gain model (i.e., akin to the hypothesis for the control observers), we simply set the pre- and post-learning gain factors to be equal; (b) For the no-conditional-inference model, we computed the estimation response on each trial as the mean of the unmodified posterior *p (**r**|s)*; (c) For the no-efficient coding model, the parameters describing the boundary-avoidance decoding template, *f, a, and w_b_,* were all set to 0, with the consequence that the likelihood *p (**r**|s)* peaked at the true stimulus motion direction and was truncated based on the implicit discrimination judgment (i.e., similar to the models in refs (Luu and Stocker, 2018; Qiu et al., 2020; Stocker and Simoncelli, 2008).

#### Tuning changes can explain human behavior nearly as well as gain modulation

We modelled the effects of training as gain modulation of sensory neurons encoding the trained stimuli, consistent with the results of (Byers et al., 2012; Byers and Serences, 2014; Chen et al., 2015). But there are other means by which encoding precision can be increased. For example, some have suggested that changes in neural tuning — i.e., changes in the density of neurons representing particular stimuli and changes in their widths — are important for PL (Adab et al., 2014; Li et al., 2004; Schoups et al., 2001; Yang and Maunsell, 2004). Along these lines, we implemented two variants of our model, in which we assumed that exposure to the trained stimuli over the learning phase of the task causes a gradual increase in the number of neurons that prefer those stimuli. A similar learning mechanism has been derived from the theory of efficient coding, in which the distributions of sensory neuron preferences are matched to the environmental distribution of stimulus attributes such as visual orientation (Ganguli and Simoncelli, 2014; Girshick et al., 2011; Wei and Stocker, 2017, 2015).

In the first model variant, we took our full model (**Figure 2A**), and simply substituted tuning changes (training-driven efficient coding) for gain modulation, leaving the (environmental) efficient coding /boundary avoidance and conditional inference steps in the original model. Specifically, we assumed that training caused neurons around the trained stimuli to change in their density, according to the solution given in ref (Ganguli and Simoncelli, 2014). These tuning changes (efficient coding) occurred such that the mutual information between the stimulus distribution and population response was maximized (Ganguli and Simoncelli, 2014). This model variant (“TC”) reproduced each aspect of human behavior well, albeit with two more parameters than the full model (**Figure 2- Figure Supplement 3**). This model had comparable goodness-of-fit and generalization performance to the full model (**Appendix 2 - Figure 1**). For nearly all observers, the optimal encoder that best explained human behavior had a higher density of neurons around the trained stimuli (±4° motion) and a lower density further away (i.e., a more exaggerated version of the warping in the pre-training population). Note that this also corresponded to higher density of neurons representing the discrimination boundary at 0°, which corresponds with a training-related/contextual prior that is higher around the boundary and trained stimuli. These model comparisons demonstrate that training-dependent tuning changes in an already non-uniform sensory population, along with implicit categorization, can explain our data as well as the original model (which assumed that training modifies sensory gain). Thus, with our current data, we cannot adjudicate between these alternatives.

In the second model variant, we tested how well such tuning changes, on their own, could account for the behavioral data. To do so, we simply removed the boundary avoidance and conditional inference components from the TC model. That is, we assumed that observers came into the experiment with a homogeneous population of sensory neurons representing motion direction, which was then modified by exposure to the training stimuli. This model captured increases in discrimination accuracy and estimation bias with learning, and their pattern of transfer across stimuli (to a lesser degree than the gain modulation model) but failed to capture the bimodality of the estimate distributions and the relations between discrimination accuracy and signed estimation accuracy. The estimation bias was under-predicted by the model and the estimation variance was over-predicted (**Figure 2- Figure Supplement 6**). Overall, this model (“TC_reduced”) had poor goodness-of-fit and generalization performance compared to the full model (**Appendix 2 - Figure 1**).

---

**Figure 1 – Figure Supplement 1.**
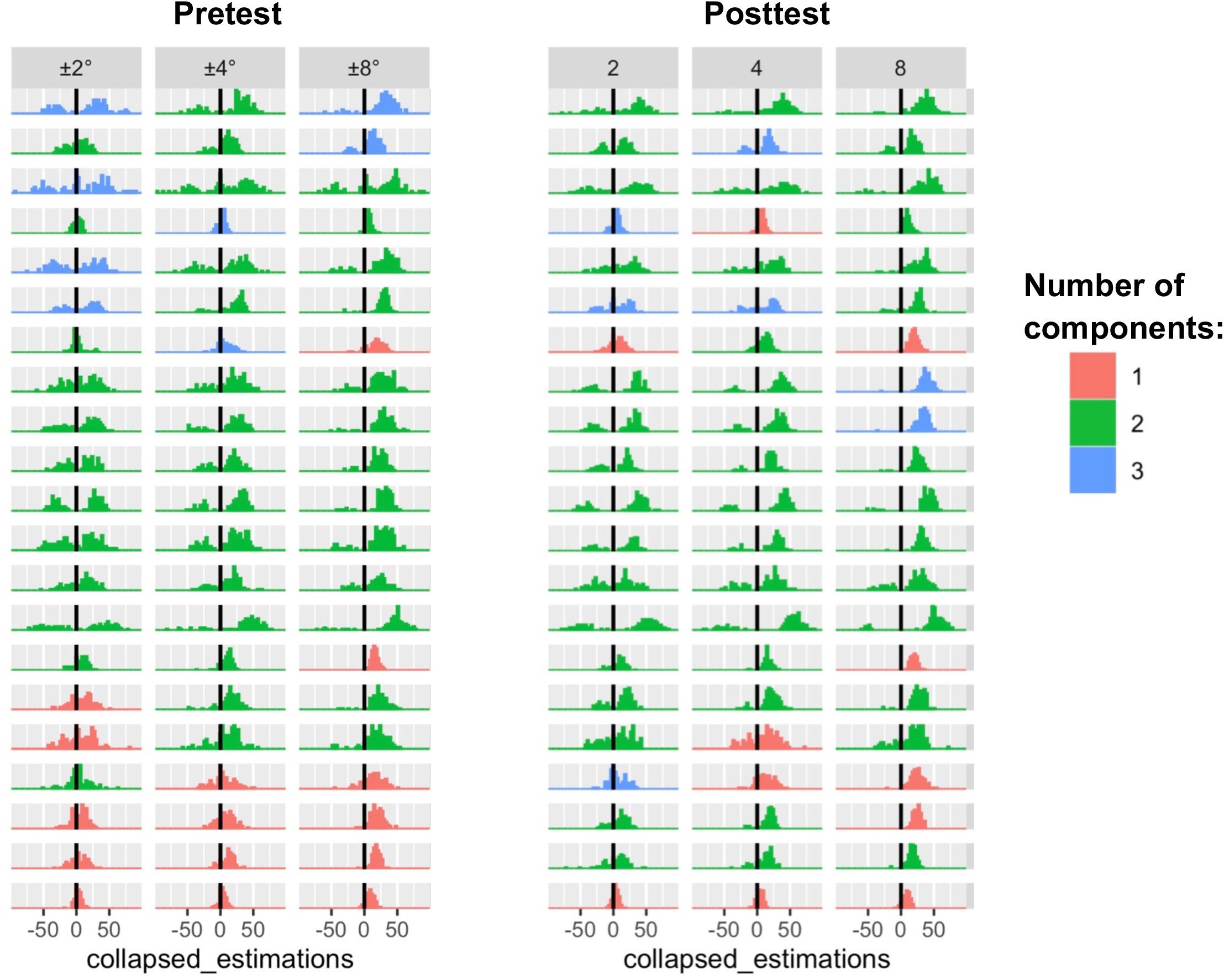
Mixture model results (used to assess level of bimodality) at pretest and posttest across all observers (rows) and directions (columns).

**Figure 1 – Figure Supplement 2.**
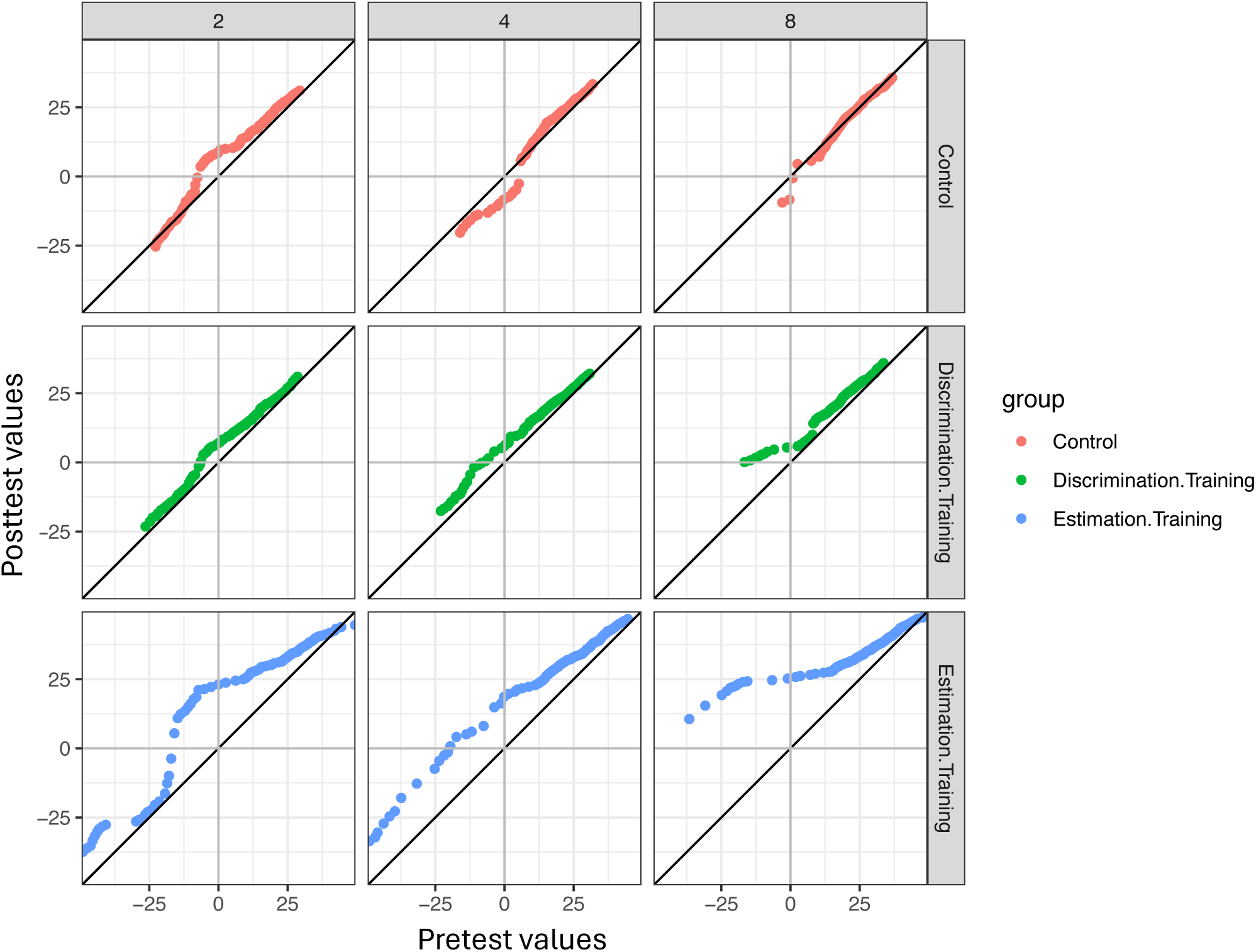
QQ-Plots of percentile values of estimate distributions at pretest versus posttest (percentile values between 0.05 and 0.95). X-axis-pretest percentile values; Y-axis-posttest percentile values. Plots represent the means across participants in each group and direction (i.e., 2°, 4°, 8°).

**Figure 1 – Figure Supplement 3.**
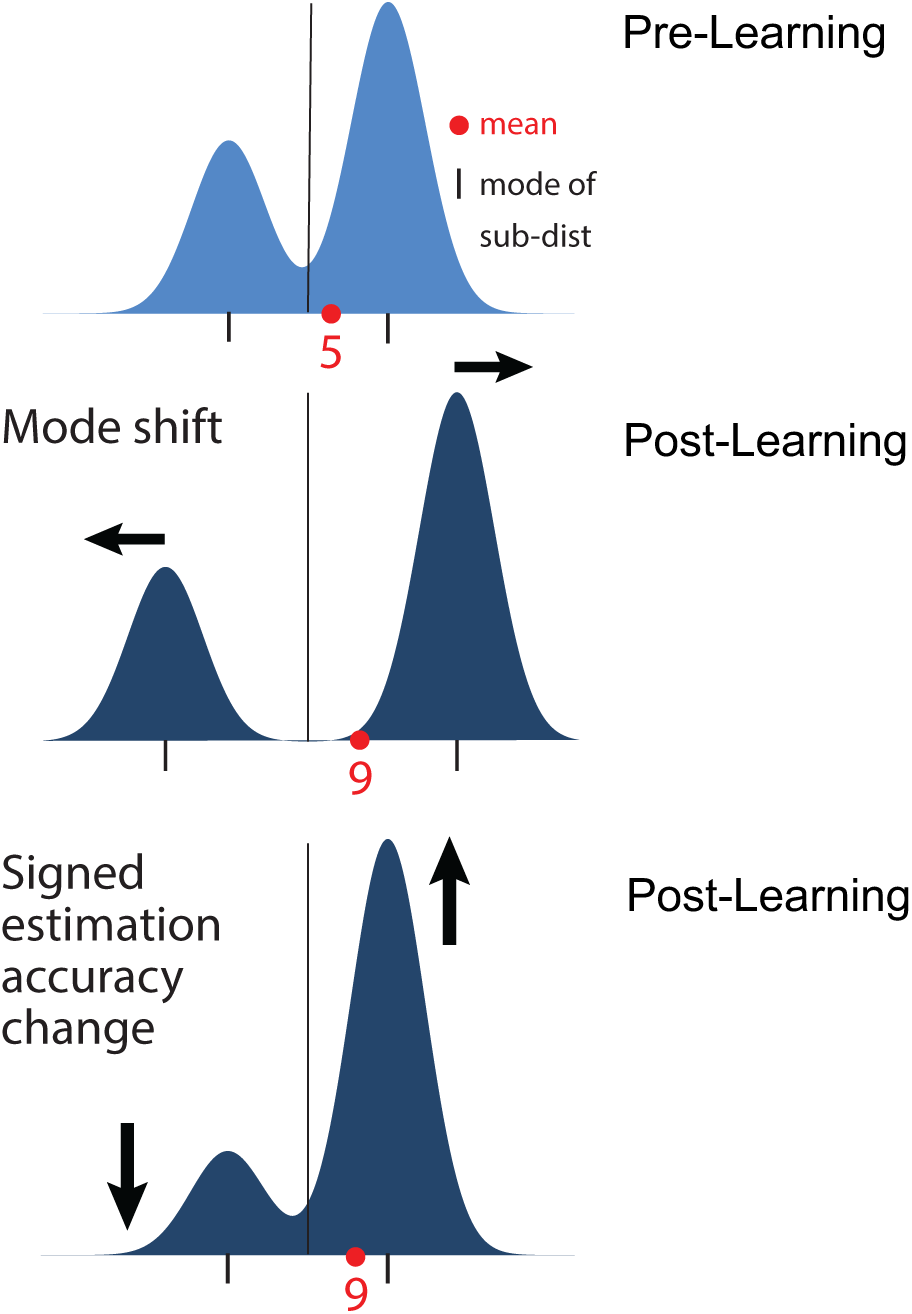
Two potential causes of overestimation. Each plot provides an idealized illustration of an estimate distribution. Top panel: pre-learning distribution, an asymmetric mixture of two Gaussian distributions. Distribution mean (red dot) lies between the two modes. Middle panel: learning-induced shift, in which the center sub-distributions are repulsed symmetrically from discrimination boundary, yielding an increase in mean, while signed estimation accuracy is unchanged. Bottom panel: learning-induced “signed estimation accuracy change”, in which the sub-distribution centers are unchanged, while their mixing proportion changes to increase signed estimation accuracy, again yielding a mean of 9°. The data provide support for a combination of these two effects.

**Figure 2 – Figure Supplement 1.**
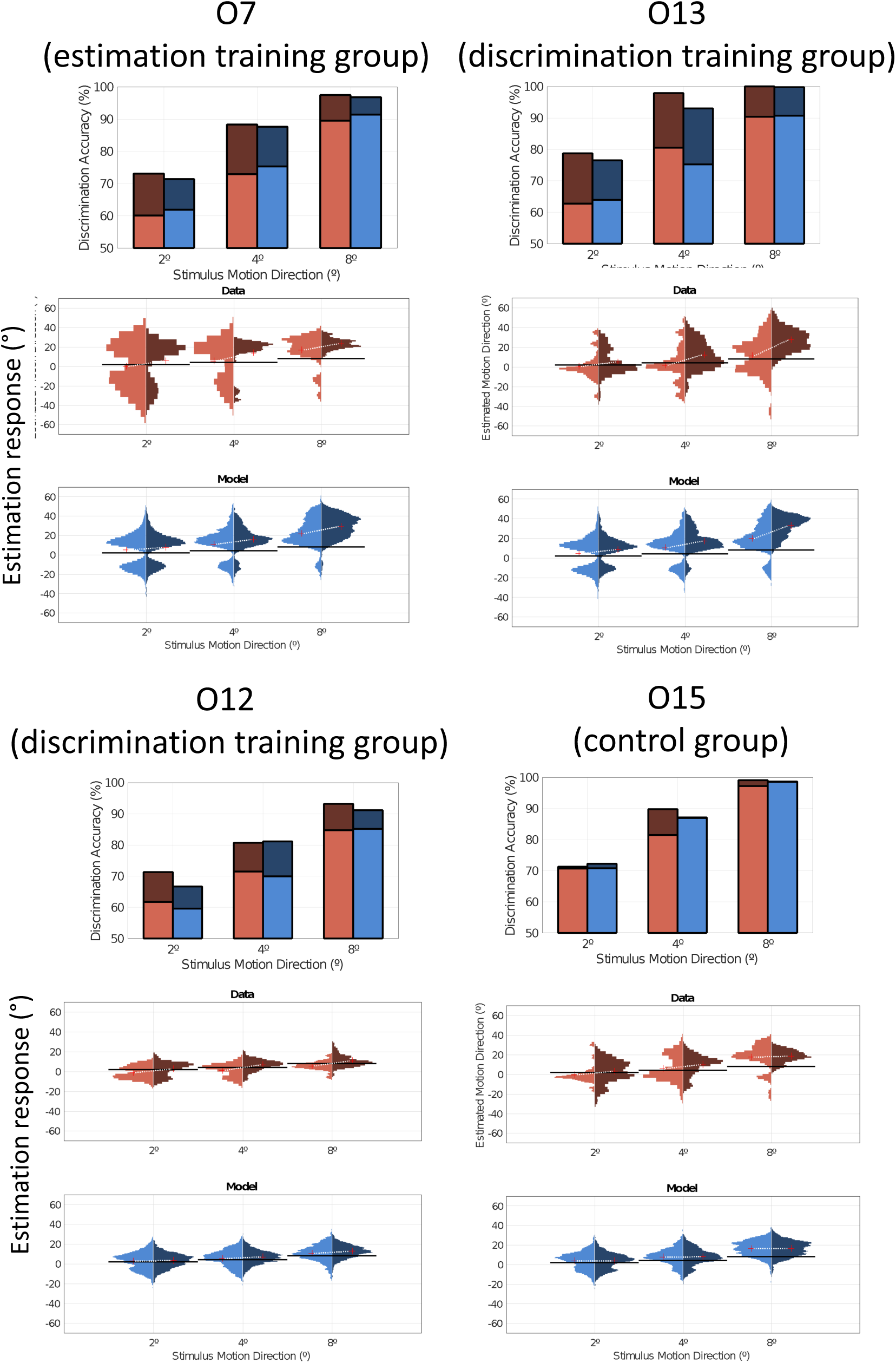
Model and Data performance. Data and model fit from representative individual observers (O7, O12, O13, O15) from each training group. Same format as Figure 2B-E.

**Figure 2 - Figure Supplement 2.**
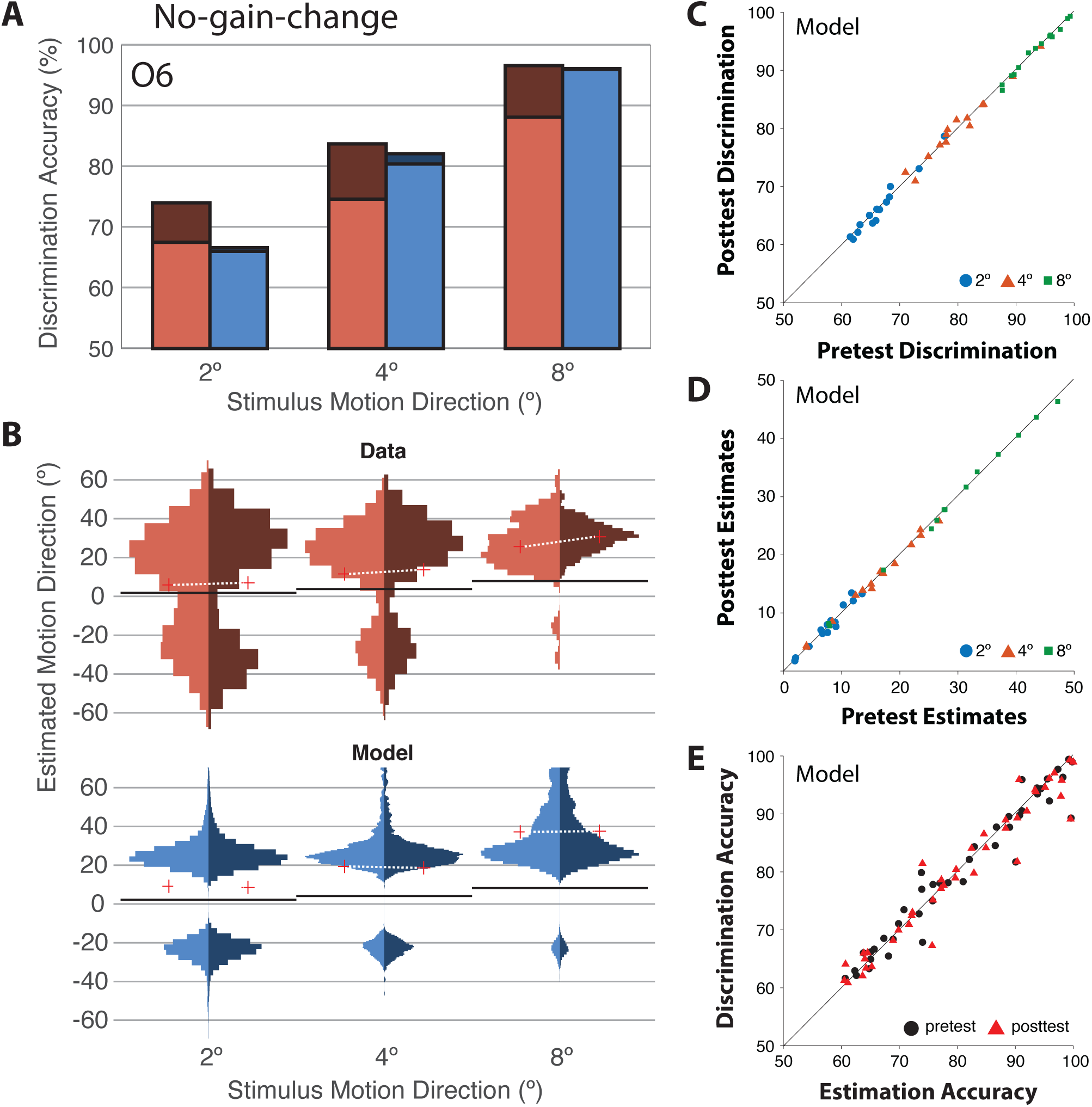
No-gain-change (“no GC”) model behavior. A variant of the full model without training-induced gain change in sensory neurons exhibits a lack of PL, but still shows a strong correlation between estimation and discrimination accuracy. Model fit to both pre- and post-training data, even though gain was fixed. For panels C-E, all trained observers included (in this and in all subsequent supplementary figures).

**Figure 2 - Figure Supplement 3.**
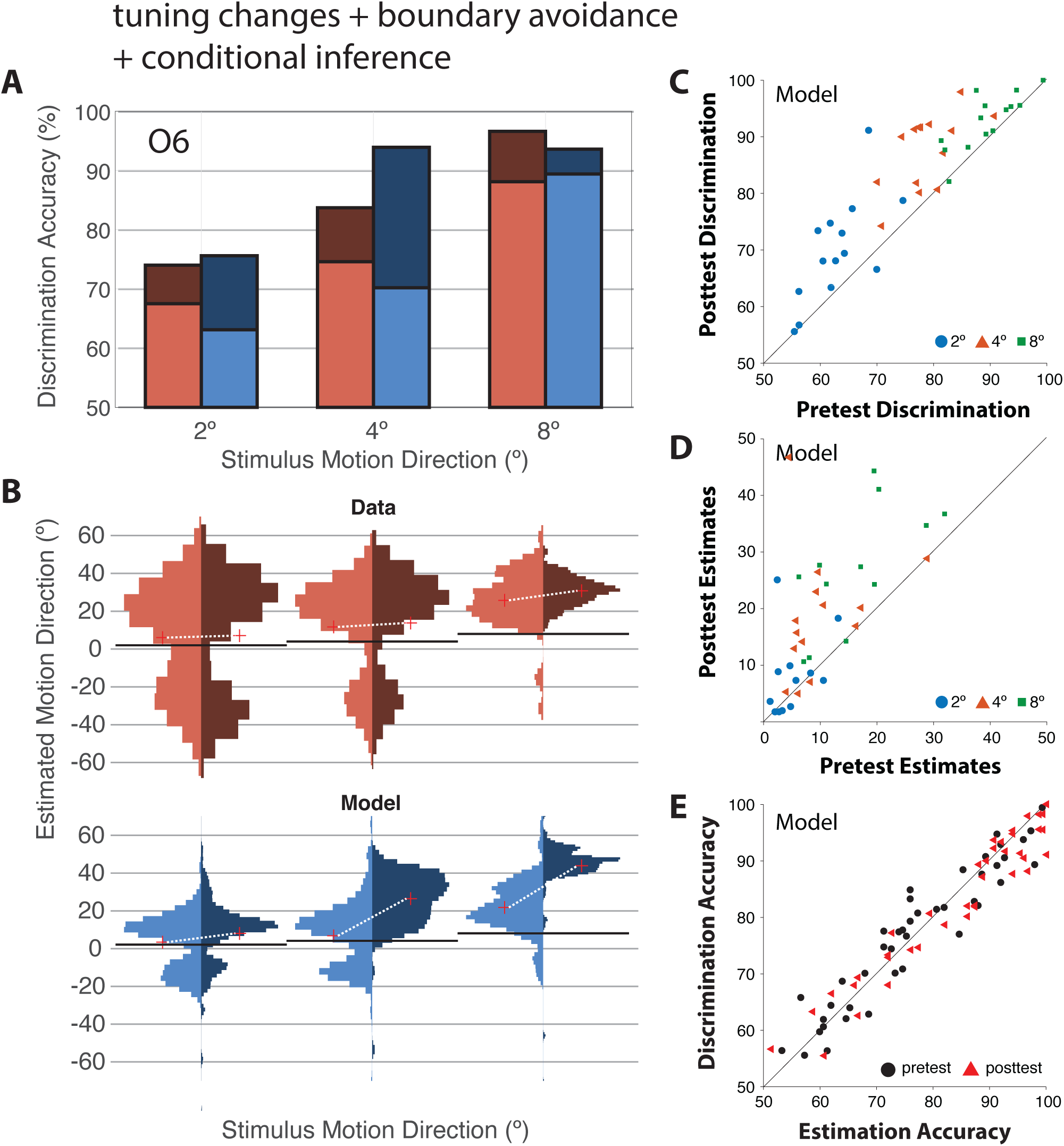
Variant of full model (gain modulation replaced with efficient coding tuning changes), “TC”. A variant of the full model, but with training-induced tuning changes instead of gain changes, shows similar behavior to the full model and fits well.

**Figure 2 - Figure Supplement 4.**
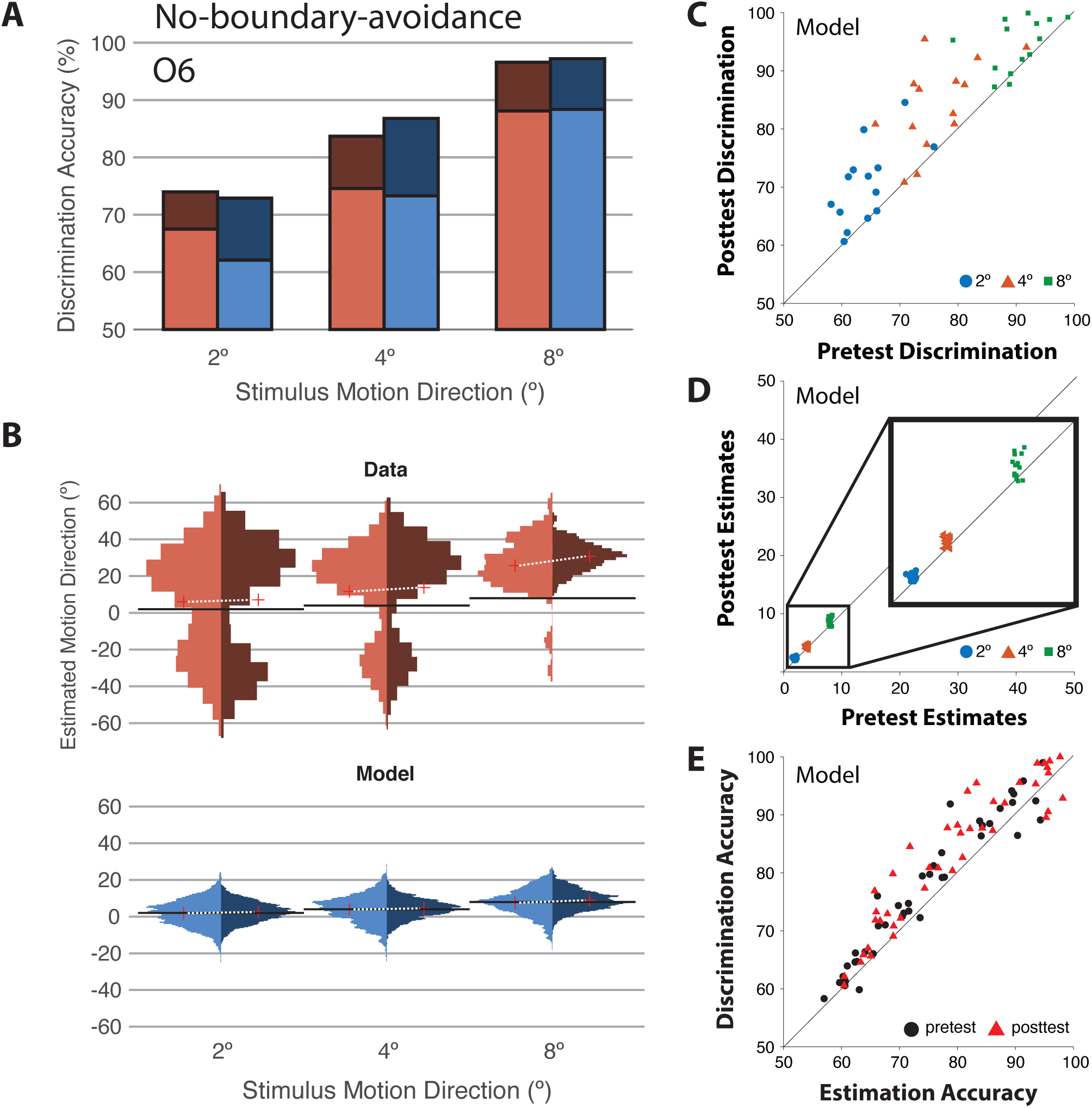
No-boundary avoidance (“no BA”) model behavior. A variant of the full model without warped tuning curves (i.e., without efficient encoding / boundary avoidance) reproduces human behavior in the discrimination task, but has much smaller biases in the estimation task (panels B and D). As a result, the relationship between signed estimation accuracy and discrimination accuracy is also different in the model than in the data, with higher discrimination accuracy than signed estimation accuracy. Inset, zoomed-in view shows that estimates are biased, but only by a few degrees.

**Figure 2 - Figure Supplement 5.**
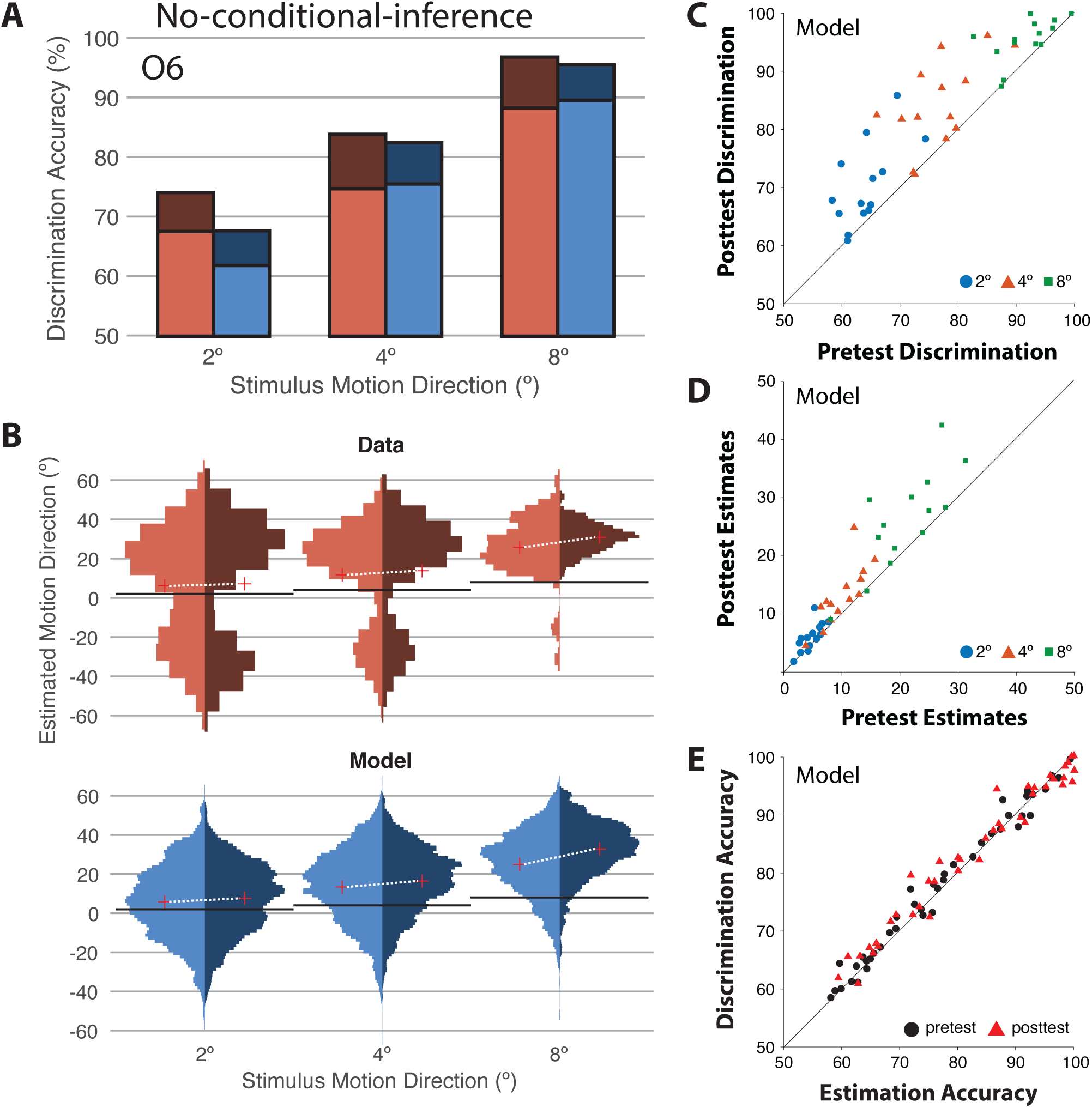
No-conditional-inference (“no CI”) model behavior. A variant of the full model without conditional inference captures human discrimination accuracy well, and captures estimation means well, but fails to capture the bimodal shape of the estimate distribution (panel B).

**Figure 2 – Figure Supplement 6.**
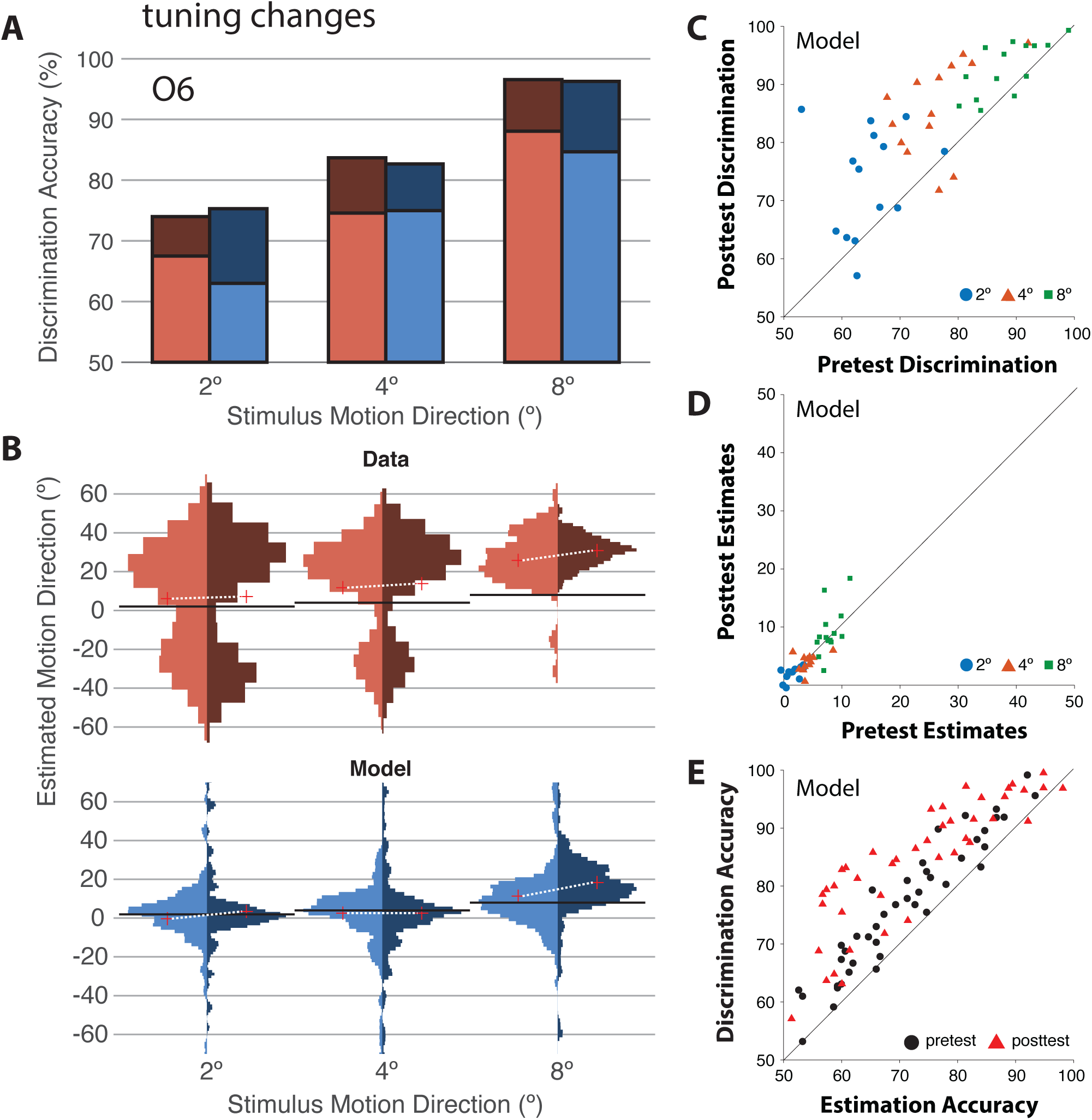
Efficient coding / tuning changes model, “TC_reduced”. A model variant in which we assumed training-induced tuning changes like in the TC model, but without boundary avoidance, gain change, and conditional inference. This model variant fails to capture the magnitude of the estimation biases and their change with learning in the data.

**Figure 2 – Figure Supplement 7.**
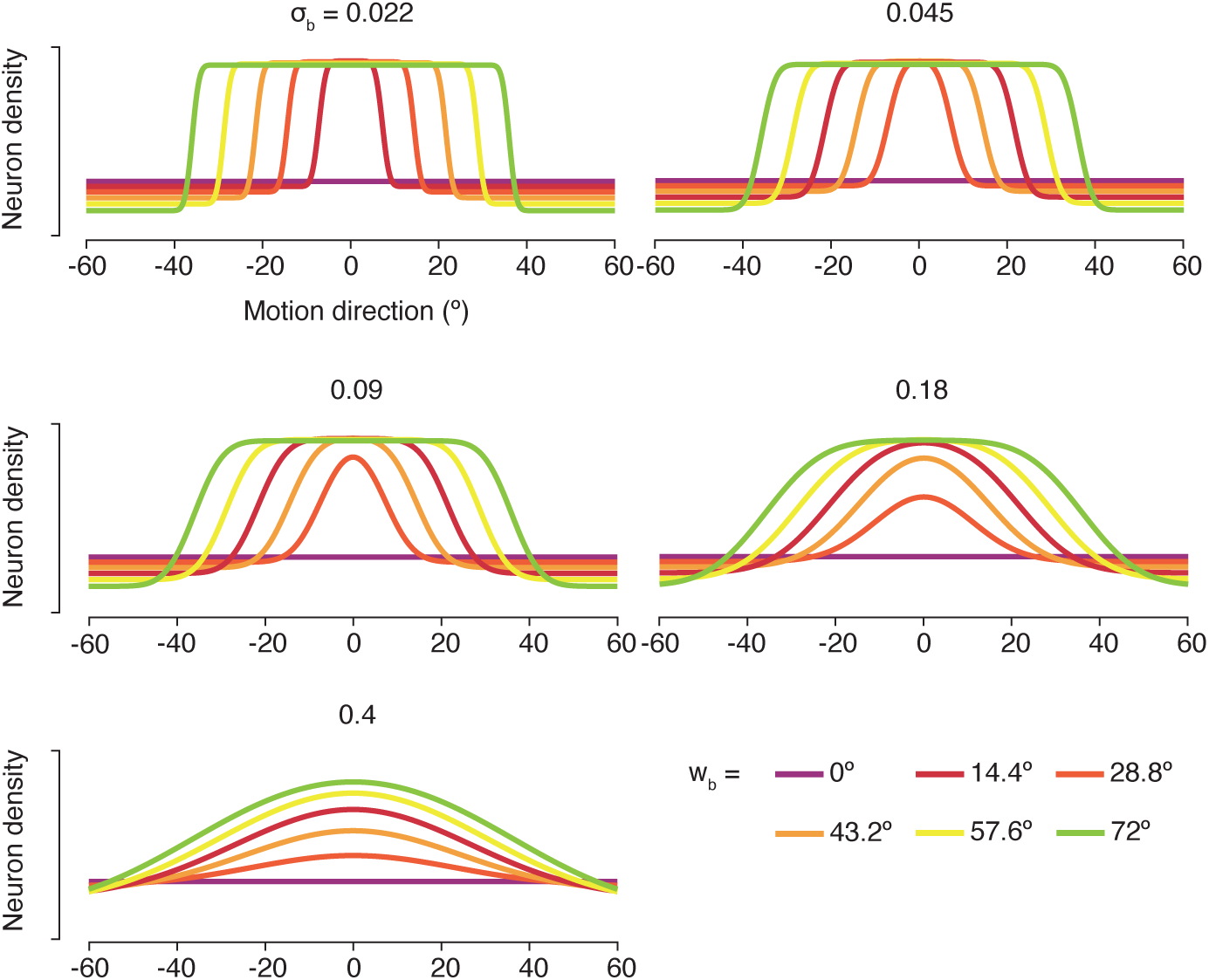
Example boundary avoidance templates, for different values of *w_b_* and *σ_b_*. Intuitively, these curves represent the extent to which the observer “avoided” the horizontal (0°) boundary in their estimation responses, and are formally defined as the derivative of the warping function that is used to re-map the sensory neurons’ tuning curves.

**Figure 3 – Figure Supplement 1.**
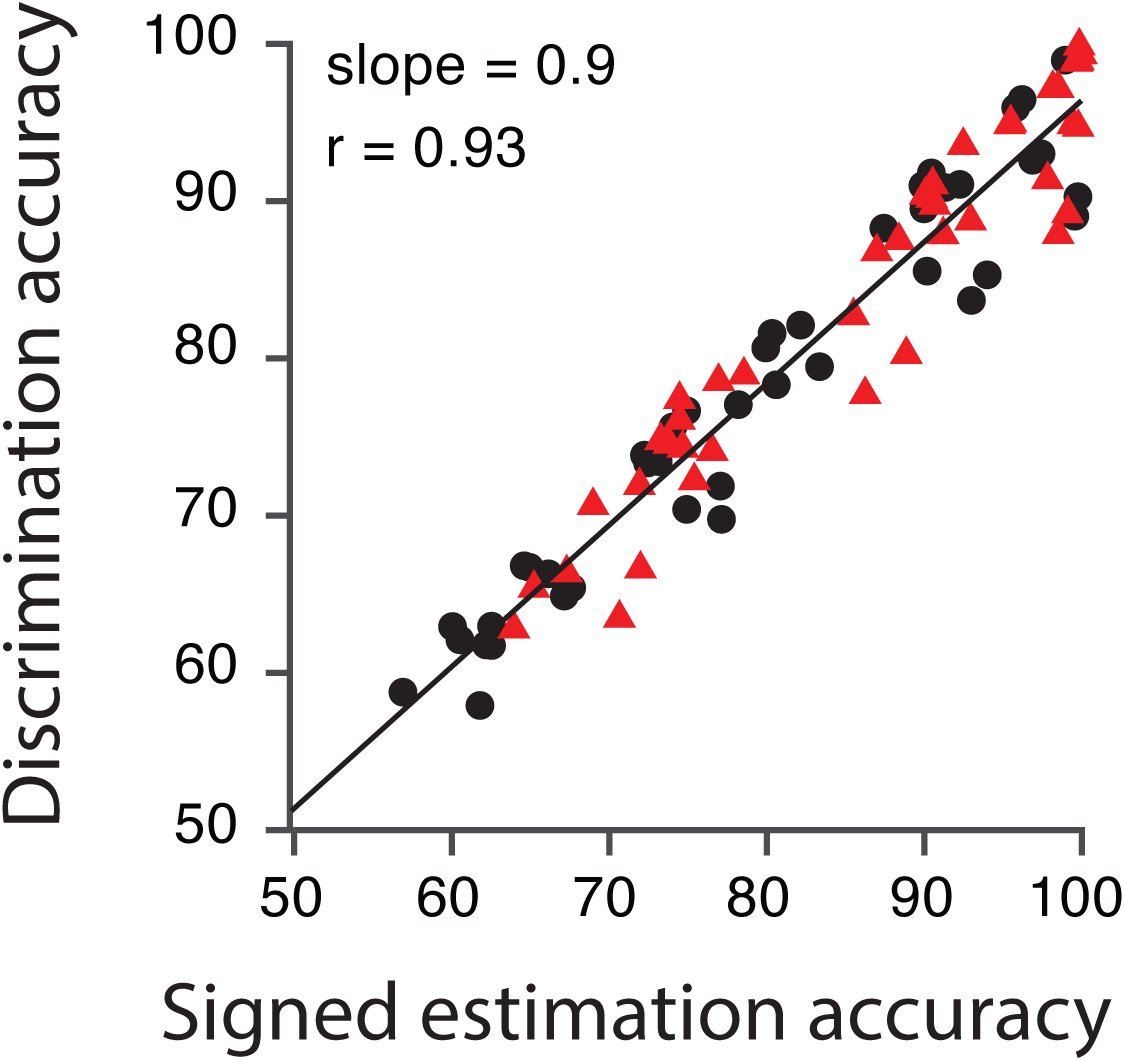
Model and Data performance. **A.** Data and model fit from representative individual observers (O7, O12, O13, O15) from each training group. Same format as Figure 2B-E. **B.** Correlation between signed estimation accuracy and discrimination accuracy in the model, collapsed across motion direction and training group for both testing sessions in pretest (black circles) and posttest (red triangles). Same format as Figure 1G.

**Figure 3 – Figure Supplement 2.**
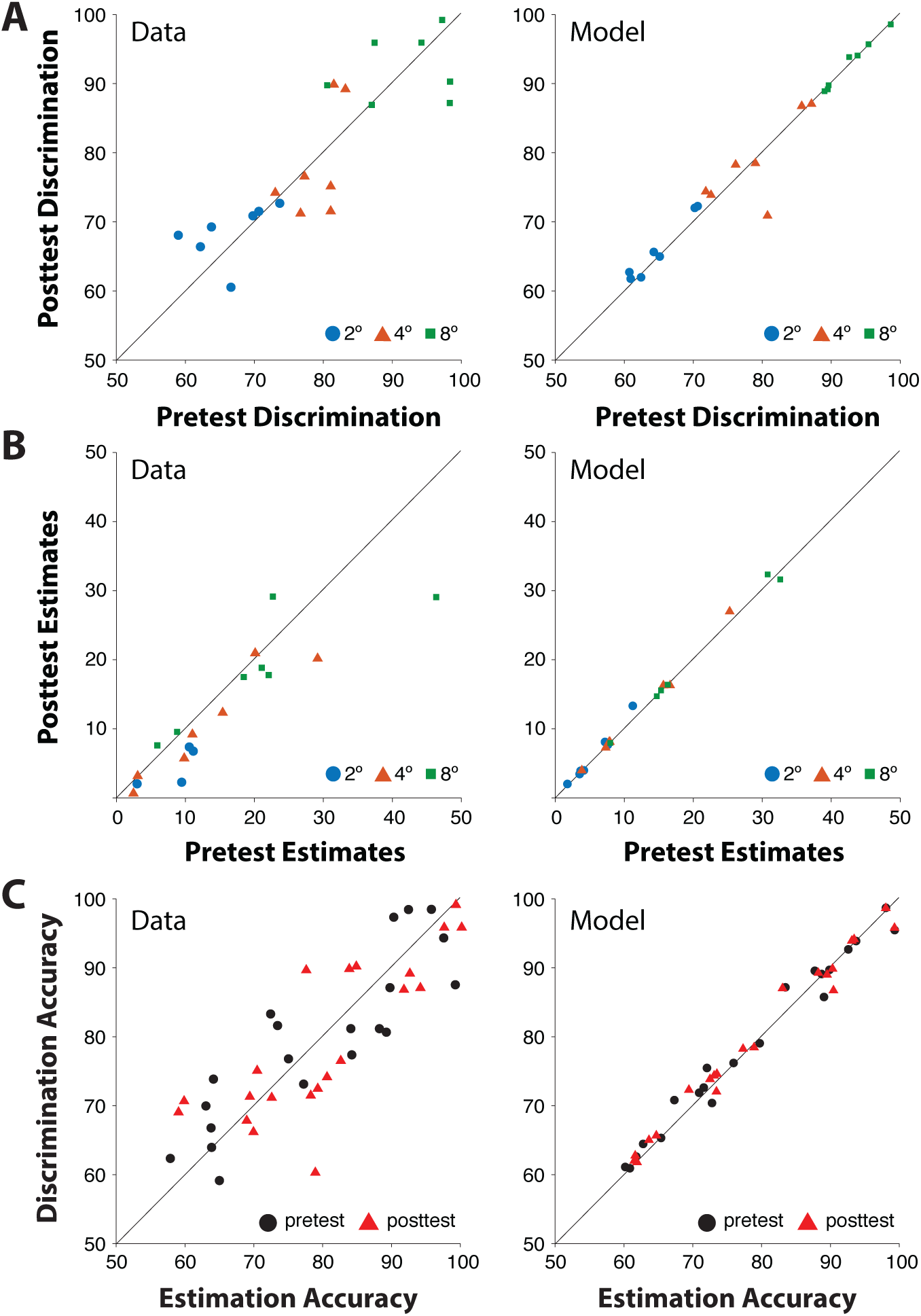
Human behavior vs model behavior for control (no training) observers. Model reproduces the lack of PL in the control group in both estimation and discrimination tasks, a correlation between estimation and discrimination accuracy, and captures inter-observer variability.

**Appendix 2 – Figure 1.**
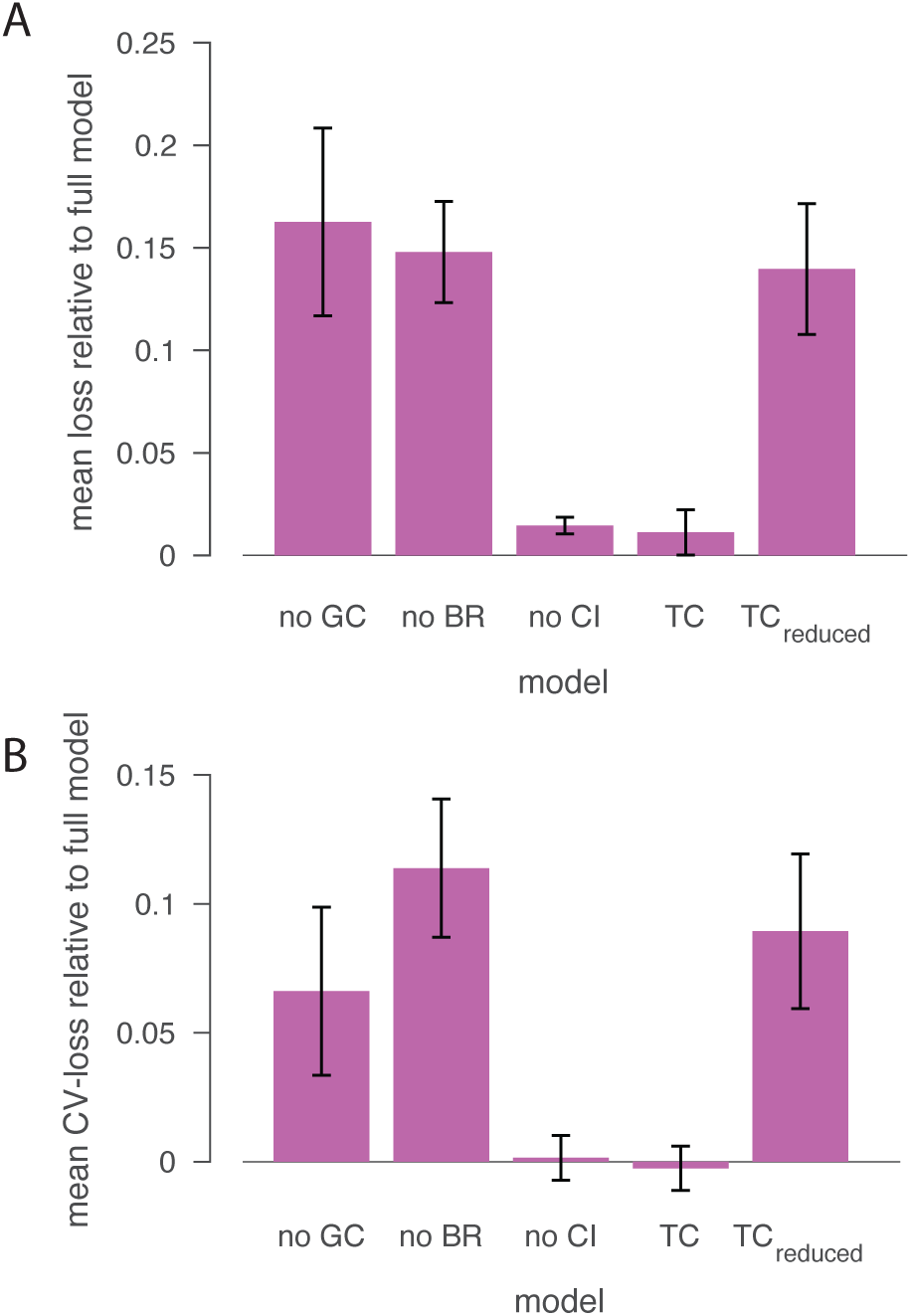
Model comparison. Models are modified versions of the full model, with: no gain change; no boundary repulsion; no conditional inference; replacement of gain changes with tuning changes (TC); and replacement of gain changes with tuning changes but without boundary repulsion or conditional inference (TC_reduced_) (Ganguli and Simoncelli, 2014, 2010). **A.** Goodness-of-fit of each model (i.e., training set / in-sample loss), averaged across iterations of model fitting (102) and observers (N = 14). Error-bars, 2 SEM across observers. **B.** Generalization performance of each model (i.e., test set / out-of-sample loss), averaged across iterations of model fitting (102) and observers (N = 14). Error-bars, 2 SEM across observers. Ultimately, we selected the full model (baseline in this figure) as the winning model.

## Notes

### Competing Interest Statement

The authors have declared no competing interest.

### Summary of Updates

Revised text and added additional analyses

